# The cell cycle controls spindle architecture in Arabidopsis by modulating the augmin pathway

**DOI:** 10.1101/2023.11.14.567058

**Authors:** Mariana Romeiro Motta, François Nédélec, Elke Woelken, Helen Saville, Claire Jacquerie, Martine Pastuglia, Sara Christina Stolze, Eveline Van De Slijke, Poyu Chen, Lev Böttger, Katia Belcram, Hirofumi Nakagami, Geert De Jaeger, David Bouchez, Arp Schnittger

## Abstract

To ensure an even segregation of chromosomes during somatic cell division, eukaryotes rely on specific microtubule structures called mitotic spindles. There are, however, striking differences in overall spindle organization among eukaryotic super groups, and in particular little is known about how spindle architecture is determined in plants. As a foundation for our work, we have measured prime characteristics of Arabidopsis mitotic spindles and built a three-dimensional dynamic model of the Arabidopsis mitotic spindle using Cytosim. Next, we identified the cell-cycle regulator CYCLIN-DEPENDENT KINASE B1 (CDKB1) together with its cyclin partner CYCB3;1 as key regulators of spindle shape and organization in Arabidopsis. Loss of CDKB1 function resulted in a high number of astral microtubules that are normally absent from plant spindles, as opposed to animal ones. We identified an augmin complex member, ENDOSPERM DEFECTIVE1 (EDE1), as a substrate of the CDKB1;1-CYCB3;1 complex. A non-phosphorylatable mutant of EDE1 displayed spindles with extended pole-to-pole distance, resembling the phenotypes of *cycb3;1* and *cdkb1* mutants. Moreover, we found that the mutated EDE1 version associated less efficiently with spindle microtubules. Consistently, reducing the level of augmin in Cytosim simulations largely recapitulated the spindle phenotypes observed in *cycb3;1* and *cdkb1* mutants. Our results emphasize the importance of cell cycle-dependent phospho-control of the mitotic spindle in plant cells. They also support the validity of our computational model as a framework for the exploration of mechanisms controlling the organization of the spindle in plants and in other species.

## Introduction

Eukaryotes have acquired specific- and robustly-functioning cytoskeletal arrays to accomplish cell divisions. Plants in particular have unique microtubule arrays for cell division, namely the preprophase band (PPB) and the phragmoplast^1^. In somatic cells, the preprophase band forms in late-G2 cells committed to division, and marks the future cortical cell-division site. After PPB disassembly and nuclear envelope breakdown, a typical barrel-shaped spindle forms, which is responsible for the segregation of sister chromatids. In telophase, the phragmoplast appears, a cytokinetic array that drives centrifugal cell plate assembly and fusion to the parental cortex. Accurate regulation of the timing and architecture of each of these microtubule structures is essential for plant morphogenesis. While the PPB and the phragmoplast have been addressed in several studies leading to important insights about their organization, relatively little is known about the mechanisms driving assembly and function of the spindle of plant cells.

Most land plants form spindles in the absence of a distinct microtubule organizing center (MTOC), responsible for nucleating microtubules in a γ-tubulin dependent manner. In animals, this MTOC is generally the centriole-containing centrosome^2^. γ-tubulin is part of the γ-tubulin ring complex (γTuRC) that acts as a template for microtubule polymerization^3^.

The augmin complex is a conserved γTuRC-targeting factor which is composed of eight members^4,5^ and allows microtubule nucleation from existing microtubules, in a parallel or branched fashion^5^. Microtubule-dependent microtubule nucleation mediated by the augmin complex amplifies microtubule number while preserving their polarity^6^. In moss, it has been shown that knocking down augmin subunits leads to a reduction of around 50% in the number of spindle microtubules^7^. Hence, augmin activity is critical for microtubule amplification and organization in the plant spindle. In Arabidopsis, ENDOSPERM DEFECTIVE1 (EDE1), an AUG8/HAUS8 homologue, targets the whole complex to spindle microtubules during mitotic cell divisions^8^. A knockdown mutant of *EDE1* displays highly elongated spindles, whereas a null mutant of this gene results in lethality^8,9^, highlighting the role of the augmin complex in plant spindle architecture.

In human cells, Polo-like kinase 1 (Plk1) has been shown to promote the association of Augmin-like complex subunit 8 (HAUS8, the human homolog of EDE1) with spindle microtubules^10^. However, plants lack Plk homologs, suggesting that cyclin-dependent kinase (CDK) complexes and/or Aurora kinases could take over some of their microtubule-associated functions in plants^11,12^. Indeed, cell-cycle factors like cyclins and CDKs are prime candidates for the regulation of spindle microtubules because of both their expression pattern as well as their localization^13,14^. In addition, plant CDK-cyclin complexes are known to be involved in the regulation of microtubule-associated proteins like MAP65-1, whose interaction with microtubules is negatively regulated by CDK phosphorylation at prophase and metaphase^15^. Thus, there is strong evidence that CDK-cyclin phosphorylation is essential for the organization and function of mitotic microtubule arrays, including the spindle^16^. Accordingly, B1-type cyclin double mutants (namely *cycb1;1 cycb1;2* and *cycb1;2 cycb1;3*) have spindles that show defects in chromosome capture, as well as other defects in the PPB and phragmoplast arrays^17^. However, little is known about the regulation of the spindle by CDK-cyclin complexes.

Here, we show that the B3-type cyclin of Arabidopsis and its main CDK partner CDKB1;1/CDKB1;2 control spindle morphogenesis. Remarkably, double *cdkb1;1 cdkb1;2* mutants displayed spindles with prominent astral microtubules reminiscent of centrosome-derived microtubules observed in animal spindles. We identify EDE1, an augmin complex member homologous to AUG8, as a substrate of the CDKB1;1-CYCB3;1 complex. Moreover, we show that a non-phosphorylatable mutant form of EDE1 results in aberrant spindle length, and this phenotype is also seen in *cycb3;1* and *cdkb1;1 cdkb1;2* mutants. Similarly, reducing augmin concentration in a 3D model of the spindle results in elongated spindles, supporting our inference of the role of cell cycle-dependent phosphorylation of augmin in plant cells.

## Results

### Generation of a computational 3D simulation of the spindle

To understand the contribution of different molecular mechanisms to the organization of the spindle, we generated a three-dimensional dynamic model of an Arabidopsis root mitotic spindle using Cytosim that extends significantly over previous simulations of the *Xenopus* spindle (Figure 1A–H and S1)^18,19^. Microtubules were generated via three different pathways: directly nucleated at the kinetochores, nucleated by augmin on the side of pre-existing microtubules, and nucleated on the spindle-poles, resulting in approximately 100, 500 and 500 microtubules in each pathway respectively. These pathways shared a cellular pool of nucleator, and microtubule assembly was limited by availability of tubulin in the cell. To simulate the spindle poles and anchor the microtubule fibers, we introduced a condensate with particle properties governed by Smoothed Particle Hydrodynamics. In addition to augmin, we included kinesin-5^20^, kinesin-14^21^, and katanin^22^ in our simulation. Kinesin-5 and kinesin-14 were added to promote microtubule cross-linking and spindle bipolarity, by sliding microtubules apart and together respectively. Notably, kinesin-5 was important to generate pulling forces on the kinetochores. Katanin was added to the condensate poles to regulate spindle length by severing. Dynein and NuMA were excluded from our simulation due to their presumed absence in plants^12^.

**Figure 1.**
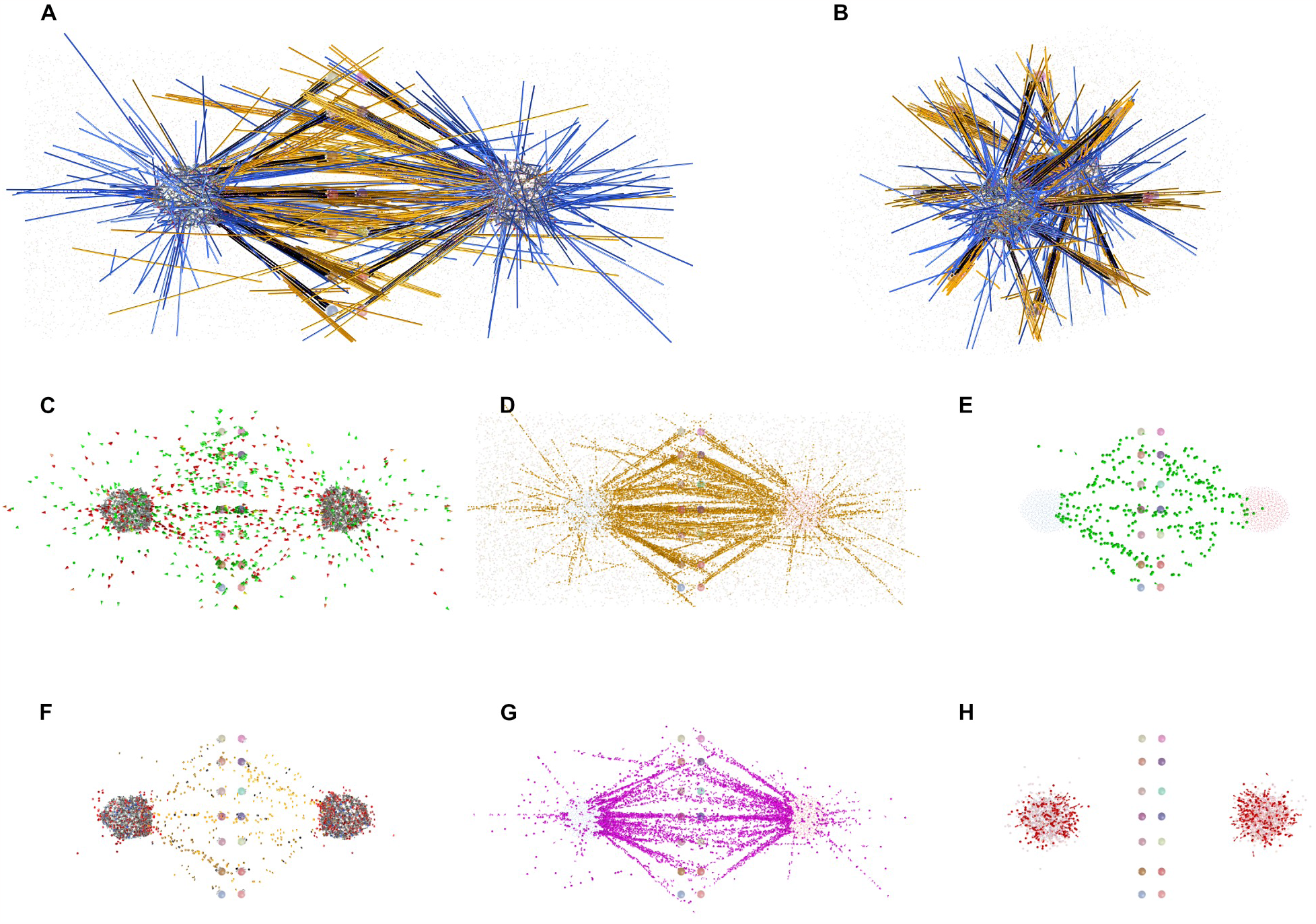
Tridimensional simulation of the Arabidopsis root mitotic spindle. (A–B) Snapshots of the simulations performed in Cytosim showing a side (A) and an end-on (B) view of the spindle. Microtubules are here color-coded according to the pathway of nucleation: blue if nucleated by the poles, black if nucleated by kinetochores, and yellow if nucleated by the augmin pathway. Chromosomes were not included in the model for simplicity, but 20 kinetochores were fixed in position such as to form a well-aligned metaphase plate. For more details of the model, see figure S1 and supplementary material. (C–H) Distribution of key elements of the simulated spindle. (C) Microtubule plus ends (red and green). (D) Kinesin-5 (yellow). (E) Augmin-activated nucleators (green). (F) Microtubule minus ends. (G) Kinesin-14 (pink). (H) Katanin (red).

Several simplifications were made, considering our focus on investigating how the general metaphase steady-state characteristics of the spindle are established. Kinetochores were fixed in position, forming a metaphase plate. When possible, spindle parameters were determined experimentally (Figure S2). First, we estimated the number of spindle microtubules by analyzing Transmission electron microscopy (TEM) images of cross-sections of Arabidopsis roots (Figure S2A and S2B). Second, the number of kinetochore microtubules was estimated by measuring the fluorescence intensity of kinetochore fibers stained against α-tubulin and by counting the number of microtubules in bundles observed by TEM (Figure S2C–G). Third, the growth rate of microtubules was measured by using a reporter fusion for the End-binding protein 1 (EB1b; Figure S2H–J)^23^. A full list of the parameters used in the simulation is provided in Table S1.

Our model produced organized spindles with focused poles and thick microtubule bundles that were attached in a bipolar manner to kinetochores (Figure 1A and 1B). At high source rates of augmin, we were able to more closely reproduce the appearance of the barrel-shaped plant spindles with few pole-nucleated microtubules (Figure 7F).

### CYCLIN B3;1 controls spindle morphogenesis

To complement our simulation approach, we sought for possible cell-cycle regulators of the plant spindle. Since we have previously shown that mitotic B1-type cyclins are key regulators of microtubule organization in Arabidopsis^17^, we decided to assess spindle shape in roots of *cycb1;1 cycb1;2* double mutants (Figure 2A). This double mutant combination has the strongest defects in growth and seed development among the B1-type cyclin mutant combinations, while still being viable^17^. We measured three spindle shape parameters, namely the lengths of major and minor axes, and the area (Figure 2B–D). Unexpectedly, the *cycb1;1 cycb1;2* mutant did not display any significant changes in spindle shape (Figure 2B–D and Table S2).

**Figure 2.**
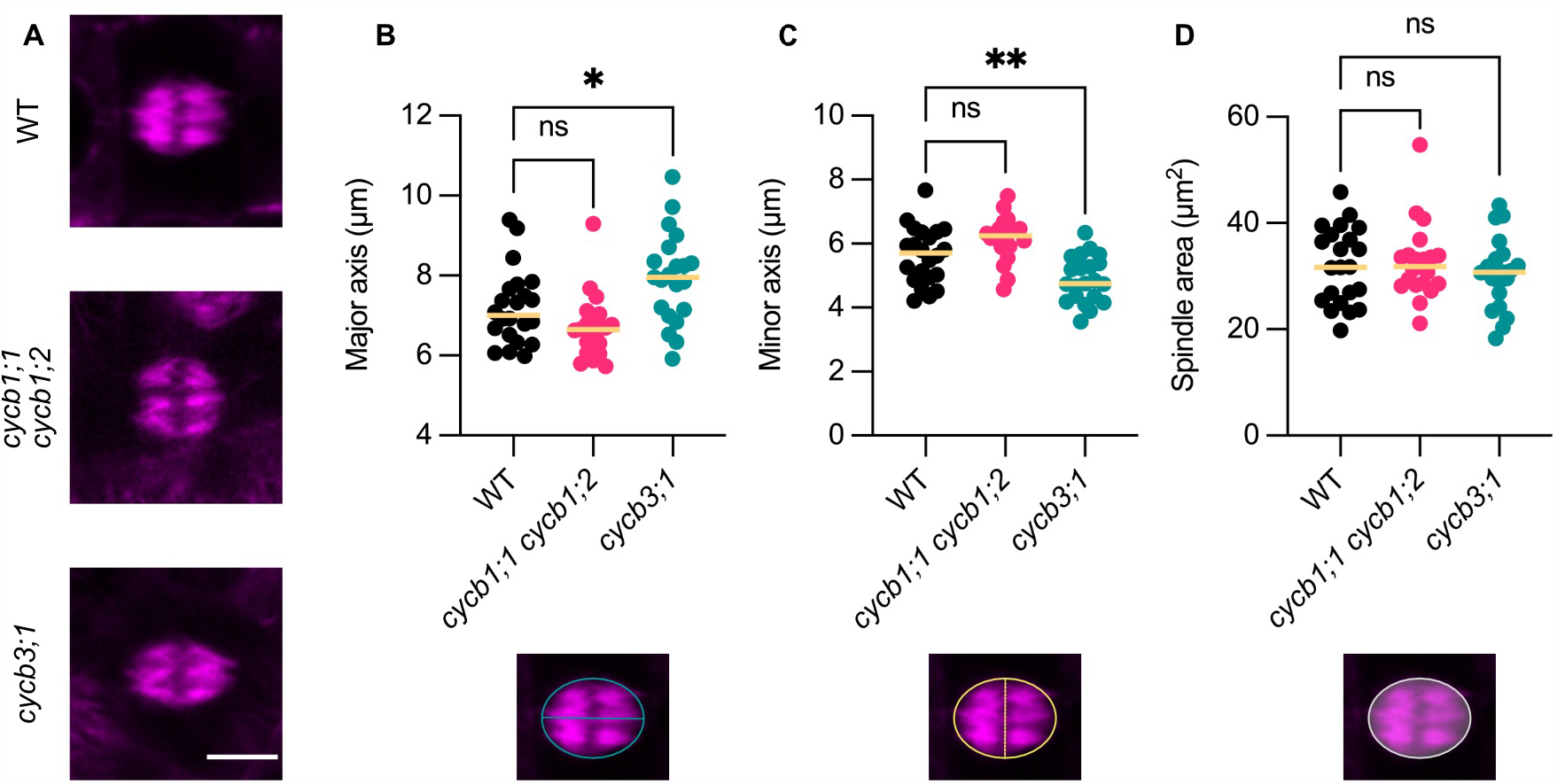
The *cycb3;1* mutant has an elongated spindle shape. (A) Confocal laser-scanning micrographs of TagRFP-TUA5-tagged microtubules in root cells at the spindle stage of WT, *cycb1;1 cycb1;2* and *cycb3;1* plants. Scale bar 5 μm. (B–D) Quantification of the spindle major axis (B), minor axis (C) and area (D) in root cells of WT (n = 22), *cycb1;1 cycb1;2* (n = 21) and *cycb3;1* (n = 21) plants. Median values were plotted as a line for each genotype. The axis or region that was measured is indicated below each graph. The level of significance was determined by an ordinary one-way ANOVA followed by Dunnett’s multiple comparisons test (* P < 0.05 and ** P < 0.01; ns depicts a non-significant difference).

We therefore hypothesized that other B-type cyclins could be involved in regulating spindle morphogenesis. The single member of the B3-type cyclin class in Arabidopsis was a good candidate as it was previously described to localize to the spindle in both mitosis and meiosis^13,14^. Indeed, spindles in roots of the *cycb3;1* mutant were more disc-shaped compared to the wild type (WT; Figure 2A) – the major axis was elongated and the minor axis was smaller, whereas the spindle area did not change significantly (Figure 2B–D and Table S2). Thus, we concluded that CYCB3;1 is a regulator of spindle morphology in Arabidopsis.

### CDKB1;1 is the main CDK partner of CYCB3;1 and the *cdkb1* mutant is hypersensitive to microtubule-destabilizing stress

To identify the main CDK partner(s) of CYCB3;1, as well as other potential interacting proteins and substrates, we performed affinity purification coupled to mass spectrometry (AP-MS) using CYCB3;1 as a bait in Arabidopsis cell suspension cultures (Figure 3B and Table S3 and S4). Five proteins were identified as potential interactors of CYCB3;1 (Figure 3B). None of them, however, were directly involved in microtubule regulation. Enzyme-substrate interactions are known to be weak and, hence, it is not surprising that we did not detect good substrate candidates in this assay. The presence of CDKB1;1 among the potential interactors, however, suggested that this kinase is the main partner of CYCB3;1. Consistently, CYCB3;1 was previously found to copurify with CDKB1;1 in tandem affinity purification experiments^24^.

**Figure 3.**
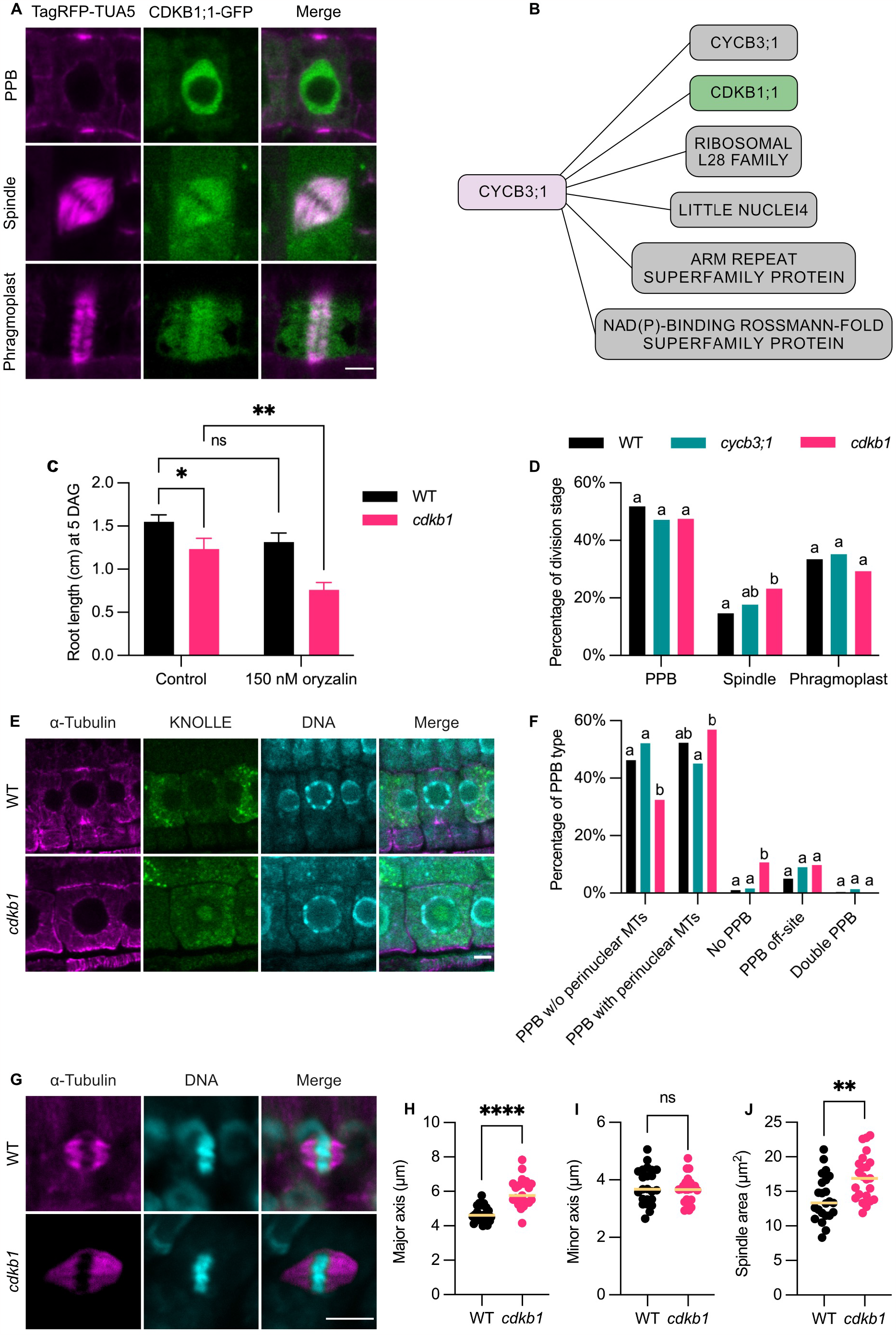
The *cdkb1* mutations affect PPB and spindle mitotic microtubule arrays. (A) Confocal laser-scanning micrographs of root cells of plants containing the TagRFP-TUA5 and CDKB1;1-GFP reporters at the three main mitotic stages (PPB, spindle and phragmoplast). The two reporters show a co-localization in the spindle and phragmoplast stages. Scale bar 5 μm. (B) Main protein interactors of CYCB3;1 as identified by AP-MS using CYCB3;1 as a bait. CDKB1;1 is highlighted in green, while other interactors that were not explored in this paper are shown in gray. (C) Quantification of root growth assays of WT and *cdkb1* seedlings on the control condition (DMSO) or 150 nM oryzalin. DAG: days after germination. Bars represent the mean value ± SD of three independent experiments with at least 16 plants per genotype per condition in each experiment. Comparisons on graph: WT control versus WT on oryzalin, *P* = 0.0843; WT control versus *cdkb1* control, *P* = 0.0211; *cdkb1* control versus *cdkb1* on oryzalin, *P* = 0.0019. (D) Quantification of PPB, spindle and phragmoplast stages in the roots of WT, *cycb3;1* and *cdkb1* plants. Different letters indicate significant differences in the proportion of the microtubule array per category in a Chi-squared test followed by the Marascuilo procedure to identify significant pairwise comparisons. Six roots were analyzed per genotype. (E) Confocal laser-scanning micrographs of cells co-stained against α-tubulin (magenta) and KNOLLE (green) in the roots of WT and *cdkb1* plants. Nuclei were counterstained with DAPI for the DNA (cyan). At this stage, the WT shows a clear accumulation of KNOLLE and a PPB, whereas the *cdkb1* mutant shows an accumulation of KNOLLE but no obvious PPB. Scale bar 5 μm. (F) Quantification of the different PPB types in the roots of WT, *cycb3;1* and *cdkb1* plants. Different letters indicate significant differences in the proportion of the PPB type per category in a Chi-squared test followed by the Marascuilo procedure to identify significant pairwise comparisons. Six roots were analyzed per genotype. (G) Confocal laser-scanning micrographs of roots cells of WT and *cdkb1* plants at the spindle stage stained against α-tubulin (magenta) and counterstained for the DNA with DAPI (cyan). Scale bar 5 μm. (H–J) Quantification of the spindle major axis (F), minor axis (G) and area (H) in the root cells of WT and *cdkb1* plants (n = 23 for both genotypes). Median values were plotted as a line for each genotype. The level of significance was determined by a two-way ANOVA followed by Tukey’s multiple comparisons test in (C) and unpaired t tests in (H–J) (* P < 0.05, ** P < 0.01, **** P < 0.0001; ns depicts a non-significant difference).

CDKB1;1 was previously shown to play a role in controlling plant growth^25^ and stomatal cell divisions^26^. CDKB1s are key regulators of DNA damage response in Arabidopsis, e.g., in response to cisplatin, by activating homologous recombination repair^27^. CDKB1s have also been shown to play a minor and partially redundant role with CDKA;1^28^, and possibly other cell-cycle kinases during cell proliferation and development of Arabidopsis. Because CDKB1;1 and CDKB1;2 have been found to function in a highly redundant manner, and likely act in similar pathways^27^, we analyzed the double mutant for these two CDKs in the following experiments.

To assess a potential role of CDKB1s in spindle regulation and track their localization in mitotic divisions, we first generated a CDKB1;1 reporter by fusing its genomic sequence to GFP. We demonstrated the functionality of the CDKB1;1-GFP reporter through its ability to rescue the root phenotype of *cdkb1* plants growing on a medium with the DNA-damaging drug cisplatin (Figure S3). In the root, the CDKB1;1-GFP reporter was found to be mainly present in the nucleus at the PPB stage, together with a faint cytosolic signal (Figure 3A). Later in mitosis, CDKB1;1-GFP co-localized with the spindle and phragmoplast microtubules (Figure 3A).

After confirming the localization of CDKB1;1 on mitotic microtubule arrays, we decided to reassess the phenotype of the *cdkb1;1 cdkb1;2* double mutant (hereafter referred to as *cdkb1*, Figure 3C–J). First, we analyzed root growth on oryzalin (Figure 3C). Oryzalin is a microtubule-destabilizing drug^29^, and many microtubule-related mutants are hypersensitive to this drug in comparison to the WT^17^. Under control conditions, the *cdkb1* mutant roots were 20.3% shorter than the WT five days after germination. Upon treatment with 150 nM oryzalin, *cdkb1* had a reduction of 38.5% in root growth, whereas, in the WT, the observed reduction in root growth was only marginally significant (Figure 3C). Thus, we concluded that the root growth phenotype of *cdkb1* is enhanced under mild microtubule destabilization conditions, prompting the hypothesis that CDKB1s could be involved in the control of mitotic microtubule arrays.

### The *cdkb1* mutant displays PPB and spindle defects

To test the role of CDKB1s in controlling microtubule organization, we first performed wholemount immunolocalization studies using antibodies against KNOLLE and α-tubulin as well as co-staining with DAPI for the DNA and counted the different mitotic stages (Figure 3D and 3F and Table S5). KNOLLE staining allows the identification of G2/M cells where PPBs are normally present in the WT^30^. First, we found that, in *cdkb1*, 10.67% of KNOLLE-positive mitotic cells had no PPB, in comparison to only 1.01% in the WT (Figure 3E and 3F), indicating that *cdkb1* mutants have defects in the establishment of the PPB. Next, we found that the *cdkb1* double mutant had a higher frequency of mitotic cells at the spindle stage in their roots (23.21%) in comparison to the WT (14.70%; Figure 3D).

We then wondered if the spindle shape of the *cdkb1* double mutants was also altered. For this analysis, we measured the spindle shape as described above in wholemount immuno-stained roots against χξ-tubulin and co-stained with DAPI (Figure 3G). Indeed, the spindles of *cdkb1* were significantly longer and larger in comparison to the WT (Figure 3G–J and Table S2). Based on these findings, we concluded that CDKB1;1 is a major regulator of mitotic microtubule arrays, particularly at the PPB and spindle stages.

### The *cycb3;1* and *cdkb1* mutants have an abnormal spindle organization and altered γ-tubulin distribution

To further characterize why the spindle shape was altered in *cycb3;1* and *cdkb1* mutants, we used super-resolution imaging with Airyscan (Figure 4). Spindles in *cdkb1* appeared highly disorganized in comparison to the WT, which could explain why they are bigger on average (Figure 3J and 4A). Furthermore, we noticed a striking number of astral microtubules in *cdkb1* spindles, which are essentially absent from the WT (Figure 4A–C). In the *cdkb1* mutant, around half of the spindles (11 out of 23 spindles) had prominent, generally short astral microtubules. This prompted us to check for the presence of astral microtubules in the *cycb3;1* mutant and, indeed, we also observed such microtubule configurations, albeit at a non-statistically significant frequency (2 out of 23 spindles; Figure 4A–C). Nevertheless, these structures were never found in the WT (n = 23).

**Figure 4.**
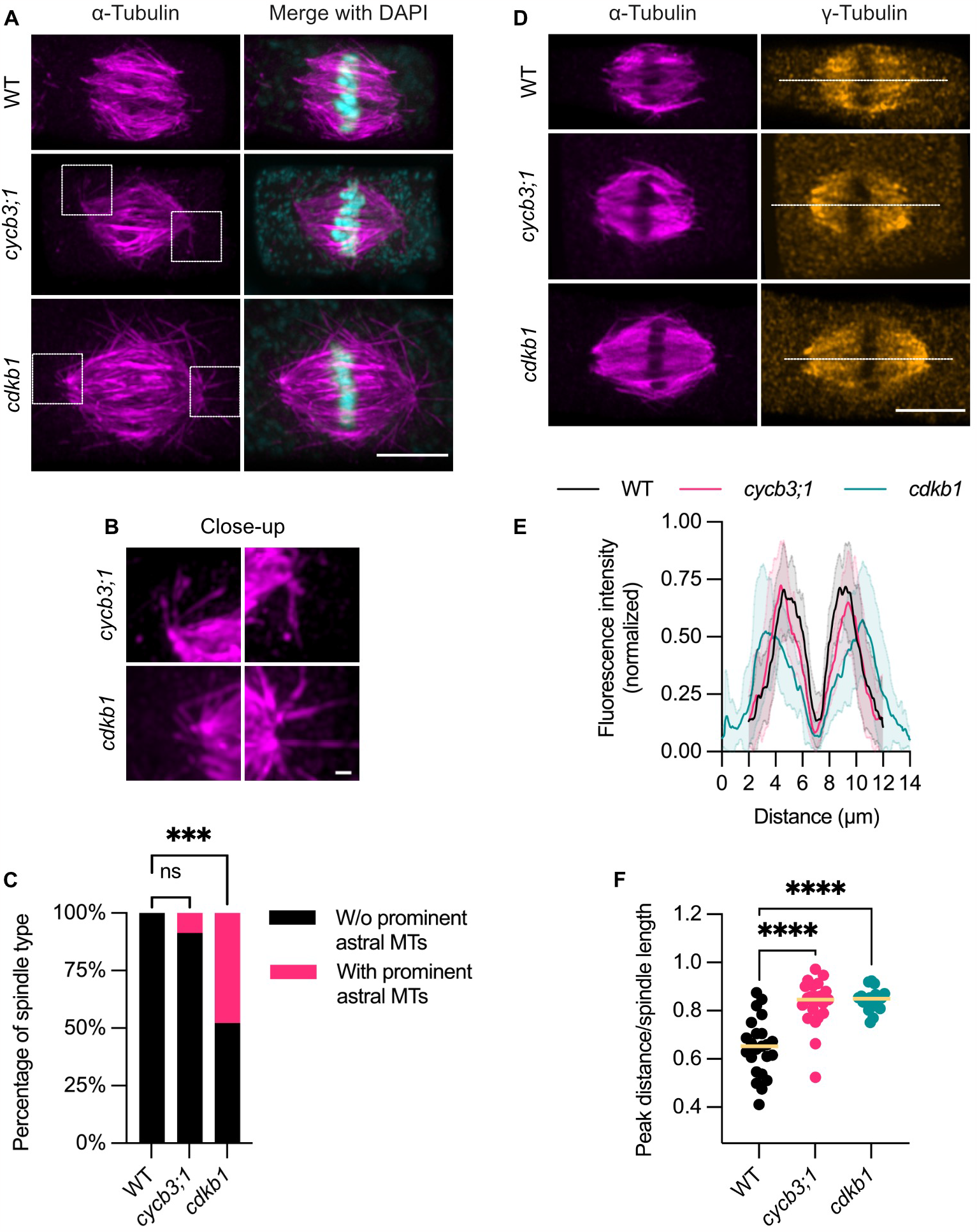
The *cycb3;1* and *cdkb1* mutants have spindles with prominent astral microtubules. (A) Maximum intensity projections of confocal laser-scanning micrographs of root cells of WT, *cycb3;1* and *cdkb1* plants at the spindle stage stained against α-tubulin (magenta) and counterstained for the DNA with DAPI (cyan). The astral microtubules are highlighted with dashed white boxes. Scale bar 5 μm. (B) Close-ups of the images shown in (A) depicting astral microtubules in the spindles of *cycb3;1* and *cdkb1* root cells stained against α-tubulin (magenta) and counterstained for the DNA with DAPI (cyan). Scale bar 0.5 μm. (C) Quantification of the number of spindles with or without prominent astral microtubules in the root cells of WT, *cycb3;1* and *cdkb1* plants (n = 23 for all genotypes). (D) Confocal laser-scanning micrographs of root cells of WT, *cycb3;1* and *cdkb1* plants at the spindle stage co-stained against α-tubulin (magenta) and γ-tubulin (orange). The white dashed line indicates the axis that was used to measure fluorescence intensity and was further plotted in the graph in (E). Scale bar 5 μm. (E) Quantification of the fluorescence intensity of γ-tubulin across the spindle axis indicated in (D) in WT (n = 23), *cycb3;1* (n = 23) and *cdkb1* (n = 22) root cells. (F) Quantification of the ratio of the distance between the fluorescence peaks seen in (E) divided by the spindle length value in WT (mean ± SD; 0.65 ± 0.12, n = 23), *cycb3;1* (0.83 ± 0.10, n = 23) and *cdkb1* (0.85 ± 0.04, n = 22) root cells. The median values were plotted as a line for each genotype. See methods for detail. The level of significance was determined by a two-proportion z-test followed by Bonferroni correction in (C) and an ordinary one-way ANOVA followed by Tukey’s multiple comparisons test in (F) (*** P < 0.001, **** P < 0.0001; ns depicts a non-significant difference).

Next, given the central function of γ-tubulin in spindle organization and function^31^, we wondered if its distribution was affected in the *cycb3;1* and *cdkb1* mutants. To that end, we performed immunostaining against χξ- and γ-tubulin in cells of the root apical meristem of the *cycb3;1* and *cdkb1* mutants (Figure 4D–F). The distribution of γ-tubulin, as expressed by the ratio of fluorescence peak distance divided by spindle length, was affected in both *cycb3;1* and *cdkb1* mutants compared to the WT (see material and methods; Figure 4F). Hence, we concluded that the localization of γ-tubulin in both *cycb3;1* and *cdkb1* mutants was strongly biased towards the spindle poles compared to the WT.

### EDE1 is a substrate of the CDKB1;1-CYCB3;1 complex and its phosphorylation is important for its function

The spindle elongation phenotype found in *cycb3;1* and *cdkb1* mutants was reminiscent of the defects previously described in *ede1* mutants^8^. EDE1 is the microtubule-binding component of the augmin complex in mitotic Arabidopsis cells. Additionally, the EDE1 protein contains eight CDK phosphorylation consensus (S-T/P) sites and was previously found to phosphorylated by human Cdk1 in *in vitro* assays^32^. Hence, we tested if the CDKB1;1-CYCB3;1 complex could phosphorylate EDE1 *in vitro*. We found that EDE1 was phosphorylated at several sites, including but not limited to at least six of the eight CDK consensus phosphorylation sites (Figure 5A and Table S6).

**Figure 5.**
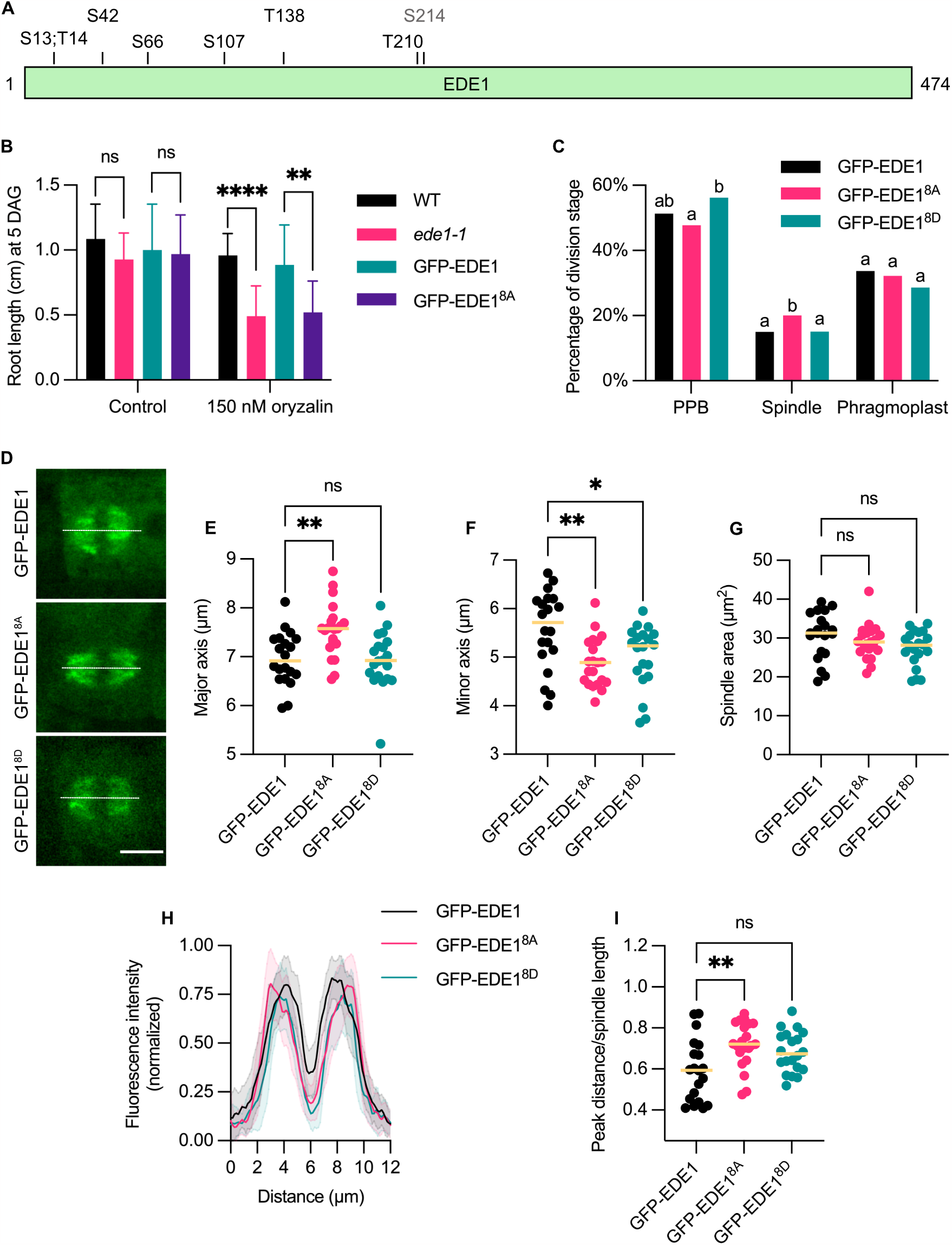
EDE1 is a substrate of the CDKB1;1-CYCB3;1 complex and its phosphorylation is important for its function. (A) Representation of the protein sequence of EDE1. All the eight mutated amino acids in the GFP-EDE1^8A^ and GFP-EDE1^8D^ constructs are represented alongside their amino acid position in the protein. Amino acids represented in black were found to be phosphorylated in the *in vitro* kinase assay with the CDKB1;1-CYCB3;1 complex, whereas the amino acid in gray (S214) was not identified in the *in vitro* kinase assay. (B) Quantification of root growth assays of WT and *ede1-1* seedlings as well as *ede1-1* mutants rescued by GFP-EDE1 or GFP-EDE1^8A^ on the control condition (DMSO) or 150 nM oryzalin. Growth on the control (mean ± SD): WT 1.08 cm ± 0.27; *ede1-1* 0.93 cm ± 0.20; *ede1*/GFP-EDE1 1.00 cm ± 0.35; and *ede1*/GFP-EDE1^8A^ 0.97 cm ± 0.30. Growth on oryzalin (mean ± SD): WT 0.96 cm ± 0.17; *ede1-1* 0.49 cm ± 0.23; *ede1*/GFP-EDE1 0.88 cm ± 0.31; and *ede1*/GFP-EDE1^8A^ 0.52 cm ± 0.24. DAG: days after germination. Bars represent the mean ± SD (n = 12–24). Two other rescue lines in the *ede1-1* background were tested for both the GFP-EDE1 and GFP-EDE1^8A^ constructs with similar results. (C) Quantification of PPB, spindle and phragmoplast stages in the roots of *ede-1* mutants rescued by GFP-EDE1, GFP-EDE1^8A^ or GFP-EDE1^8D^. Different letters indicate significant differences in the proportion of the microtubule array per category in a Chi-squared test followed by the Marascuilo procedure to identify significant pairwise comparisons. Seven roots were analyzed per genotype. (D) Confocal laser-scanning micrographs of GFP-EDE1-tagged microtubules in root cells at the spindle stage of *ede1-1* mutants rescued by GFP-EDE1, GFP-EDE1^8A^ or GFP-EDE1^8D^. Scale bar 5 μm. (E–G) Quantification of the spindle major axis (E), minor axis (F) and area (G) in the root cells of *ede1-1* mutants rescued by GFP-EDE1 (n = 20), GFP-EDE1^8A^ (n = 21) or GFP-EDE1^8D^ (n = 20). Median values were plotted as a line for each genotype. (H) Quantification of the fluorescence intensity of GFP-EDE1 across the spindle axis indicated in (D) in root cells of *ede1-1* mutants rescued by GFP-EDE1 (n = 20), GFP-EDE1^8A^ (n = 21) or GFP-EDE1^8D^ (n = 20). (I) Quantification of the ratio of the distance between the fluorescence peaks seen in (H) divided by the spindle length value in root cells of *ede1-1* mutants rescued by GFP-EDE1 (n = 20), GFP-EDE1^8A^ (n = 21) and GFP-EDE1^8D^ (n = 20). The median values were plotted as a line for each genotype. Comparisons on graph: GFP-EDE1 versus GFP-EDE1^8A^, *P* = 0.0048; GFP-EDE1 versus GFP-EDE1^8D^, *P* = 0.0610. See methods for detail. The level of significance was determined by a two-way ANOVA followed by Tukey’s multiple comparisons test in (B) and one-way ANOVAs followed by Tukey’s multiple comparisons tests in (E–G) and (I) (* P < 0.05, ** P < 0.01, **** P < 0.0001; ns depicts a non-significant difference).

To address the localization of EDE1 in mitosis and assess the importance of its phosphorylation, we first generated a genomic EDE1 reporter (GFP-EDE1). We also mutated eight CDK phosphosites (seven of them identified *in vitro*) into either an alanine (GFP-EDE1^8A^), which blocks phosphorylation, or an aspartate (GFP-EDE1^8D^), which mimics a phosphorylated amino acid (Figure 5A). We introduced the WT and mutated versions in the *ede1-1* mutant background (hereafter referred to as *ede1*/GFP-EDE1, *ede1*/GFP-EDE1^8A^ and *ede1/*GFP-EDE1^8D^). The *ede1*/GFP-EDE1^8A^ plants had a fully rescued seed phenotype (Figure S4A and S4B). However, we found that their root growth was hypersensitive to oryzalin, similarly to the *ede1-1* mutant, whereas *ede1*/GFP-EDE1 plants grew similarly to the WT (Figure 5B). When we measured the timing from nuclear envelope breakdown (NEB) to anaphase onset (AO) with or without 150 nM oryzalin in *ede1*/GFP-EDE1 plants, we did not find a significant change (Figure 6A and 6B). In contrast, in *ede1*/GFP-EDE1^8A^ plants, the NEB to AO duration was significantly longer in oryzalin-treated plants (Figure 6A and 6B). This showed that the functionality of the non-phosphorylatable GFP-EDE1^8A^ protein was affected, especially under stress conditions.

**Figure 6.**
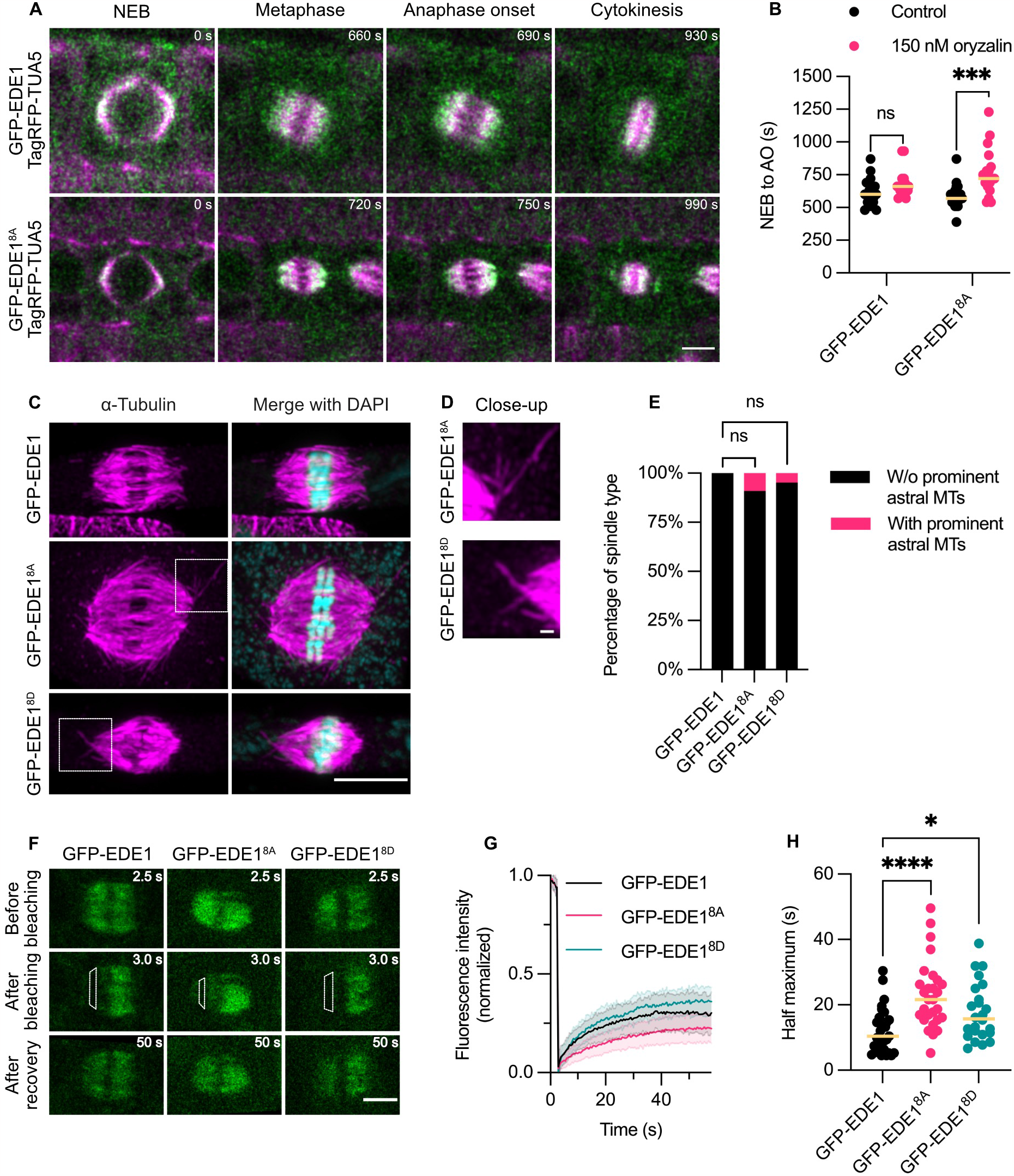
The phosphorylation of EDE1 is important for its localization at spindle microtubules. (A) Confocal laser-scanning micrographs of GFP-EDE1- and TagRFP-TUA5-tagged microtubules in root cells of *ede1-1* mutants rescued by GFP-EDE1 and GFP-EDE1^8A^. Mitotic cells were followed from nuclear envelope breakdown (NEB) through the anaphase onset stage (AO) to cytokinesis. The timepoint is indicated on the top right of the images in seconds. Scale bar 5 μm. (B) Quantification of the length of the NEB to AO stage in root cells of *ede1-1* mutants rescued by GFP-EDE1 and GFP-EDE1^8A^ on the control (mean ± SD; 617.6 s ± 104.0 for GFP-EDE1 and 588.3 s ± 98.8 for GFP-EDE1^8A^, n = 17–18) or 150 nM oryzalin condition (681.2 s ± 102.7 for GFP-EDE1 and 765.0 s ± 180.7 for GFP-EDE1^8A^, n = 17–18). The median values were plotted as a line for each genotype and condition. Comparisons on graph: GFP-EDE1 control versus GFP-EDE1 on oryzalin, *P* = 0.4673; GFP-EDE1^8A^ control versus GFP-EDE1^8A^ on oryzalin, *P* = 0.0005. (C) Maximum intensity projections of confocal laser-scanning micrographs of root cells of *ede1-1* mutants rescued by GFP-EDE1, GFP-EDE1^8A^ and GFP-EDE1^8D^ at the spindle stage stained against α-tubulin (magenta) and counterstained for the DNA with DAPI (cyan). The astral microtubules are highlighted with dashed white boxes. Scale bar 5 μm. (D) Close-ups of the images shown in (C) depicting astral microtubules in the spindles of *ede1-1* mutant root cells rescued by GFP-EDE1^8A^ and GFP-EDE1^8D^ and stained against α-tubulin (magenta) and counterstained for the DNA with DAPI (cyan). Scale bar 0.5 μm. (E) Quantification of spindles with or without prominent astral microtubules in the root cells of *ede1-1* mutants rescued by GFP-EDE1 (n = 12), GFP-EDE1^8A^ (n = 22) and GFP-EDE1^8D^ (n = 21). (F) Confocal laser-scanning micrographs of root cells in which the FRAP assay of spindles tagged by GFP-EDE1, GFP-EDE1^8A^ or GFP-EDE1^8D^ in the *ede1-1* background was performed. The white dashed box represents the area that was bleached by the laser. The time is indicated on the top right of the images in seconds. Scale bar 5 μm. (G) Quantification of the fluorescence intensity recovery over time following bleaching of spindles in root cells tagged by GFP-EDE1 (n = 31), GFP-EDE1^8A^ (n = 28) or GFP-EDE1^8D^ (n = 24) in the *ede1-1* background. The fluorescence intensity was normalized in each cell by the maximum and minimum values and plotted as an average (line) ± SD (shaded area). (H) Quantification of the half maximum values in seconds of fluorescence recovery in *ede1-1* mutants rescued by GFP-EDE1 (n = 31), GFP-EDE1^8A^ (n = 28) or GFP-EDE1^8D^ (n = 24). The median values were plotted as a line for each genotype. Comparisons on graph: GFP-EDE1 versus GFP-EDE1^8A^, *P* < 0.0001; GFP-EDE1 versus GFP-EDE1^8D^, *P* = 0.0452. The level of significance was determined by ordinary one-way ANOVAs followed by Tukey’s multiple comparisons tests in (B) and (H) and a two-proportion z-test followed by Bonferroni correction in (E) (* P < 0.05, *** P < 0.001, **** P < 0.0001; ns depicts a non-significant difference).

To further characterize mitotic defects in *ede1-1* plants rescued by the different GFP-EDE1 versions, we measured the frequency of PPB, spindle and phragmoplast stages in root apical meristems (Figure 5C and Table S5). Similar to *cdkb1* mutants, *ede1*/GFP-EDE1^8A^ had a significant overrepresentation of spindle stages in mitotic cells (20.05% of total mitotic figures versus 14.99% in *ede1*/GFP-EDE1). We found that *ede1*/GFP-EDE1^8A^ plants displayed deformed spindles highly reminiscent of *cycb3;1* (Figure 5D–G). Their major axis was larger and their minor axis was smaller in comparison to *ede1*/GFP-EDE1, whereas the spindle area did not change significantly (Table S2). Conversely, *ede1*/GFP-EDE1^8D^ did not have a significant change in the major axis or spindle area compared to *ede1*/GFP-EDE1, but still had a significantly smaller minor axis, albeit not as reduced as in *ede1*/GFP-EDE1^8A^ (Table S2). We concluded that EDE1 phosphorylation has an impact on spindle architecture under control conditions, and becomes even more critical when microtubules are destabilized.

Based on the striking similarity between the phenotypes of *cycb3;1* and *ede1*/GFP-EDE1^8A^, we hypothesized that EDE1 is a major substrate of CDK-cyclin complexes involving CYCB3;1. To test this hypothesis, we made crosses of *cycb3;1* with *ede1-1* mutants. Indeed, spindle defects in *cycb3;1 ede1-1* double mutants were identical to the single *ede1-1* mutant (Figure S5A–D). We thus concluded that EDE1 is the main substrate of CYCB3;1 action, whereas the *cdkb1* mutant phenotype is possibly more pleiotropic and a result of alterations in different CDK-cyclin phosphorylation pathways.

### EDE1 phosphorylation is important for its localization at the spindle

Since the human homolog of EDE1 has been suggested to stabilize microtubules^33^, we wondered if *ede1*/GFP-EDE1^8A^ plants had impaired tubulin turnover^34^, which results from the combination of many microtubule activities including dynamic instability and could contribute to the above-described phenotypes. To test that, we performed a FRAP assay of microtubules tagged with TagRFP-TUA5 in the *ede1*/GFP-EDE1 or *ede1*/GFP-EDE1^8A^ backgrounds and observed their recovery over time (Figure S4C–E). However, we did not find a significant difference in the half maximum values between the two genotypes and fluorescence recovered at similar rates in both cases. Thus, we concluded that tubulin turnover did not change significantly in *ede1*/GFP-EDE1^8A^ plants in comparison to *ede1*/GFP-EDE1.

As EDE1 is known to recruit the γTuRC to spindle microtubules, and given the biased distribution of γ-tubulin in the *cycb3;1* and *cdkb1* mutants, we assessed the localization of the mutated forms of GFP-EDE1 at the spindle in the *ede1-1* background (Figure 5D, 5H and 5I). Indeed, the distribution of GFP-EDE1^8A^ was significantly biased towards the spindle poles in comparison to GFP-EDE1, as expressed by the ratio of peak distance divided by spindle length, whereas the GFP-EDE1^8D^ version did not show a significant difference in localization in comparison to GFP-EDE1 (Figure 5H and 5I). In addition, we found that spindles of *ede1/*GFP-EDE1^8A^ plants also displayed prominent astral microtubules in 2 out of 22 cases (Figure 6C–E), reminiscent of the *cycb3;1* mutant (2 out of 23 spindles). Spindles of *ede1*/GFP-EDE1^8D^ plants also displayed astral microtubules, although at a lower frequency (1 out of 21 spindles). Though the differences were not statistically significant regarding the proportion of spindles displaying astral microtubules in *ede1*/GFP-EDE1^8A^ or *ede1*/GFP-EDE1^8D^ in comparison to *ede1*/GFP-EDE1 (Figure 6E), we have shown above that these structures were never found in wild-type spindles (n = 23) and, accordingly, no prominent astral microtubules were found in *ede1*/GFP-EDE1 (n = 12).

Since the binding of HAUS8 to microtubules is enhanced upon phosphorylation by Plk1^10^, we wondered if the phosphorylation of EDE1 also affected its association with microtubules. We thus performed FRAP assays in spindles of *ede1*/GFP-EDE1, *ede1*/GFP-EDE1^8A^ and *ede1*/GFP-EDE1^8D^ root cells (Figure 6F–H). The half maximum of GFP-EDE1^8A^ was on average 22.96 s ± 10.42, significantly longer than GFP-EDE1 (12.04 s ± 6.72). GFP-EDE1^8D^ had an average half maximum of 17.79 s ± 8.78, further confirming that it functions more similarly to GFP-EDE1 than the GFP-EDE1^8A^ version, although this was still a significantly slower recovery compared to GFP-EDE1. Therefore, we concluded that the phosphorylation of EDE1 is important for its association with spindle microtubules and is significantly blocked in the GFP-EDE1^8A^ protein.

### Altering the amount of augmin in the simulation affects spindle length and organization

To validate the role of augmin in overall spindle organization, and considering our experimental observations, we manipulated the amount of augmin in our simulations (Figure 7A–I, n = 132 simulations of 2000 s). As the augmin source rate increased (while all other parameters were constant), spindle length decreased – quickly at first and then slowly (Figure 7A). In the range of augmin source rates we tested, the spindle length decreased by about 50%. With increasing augmin source rates, the average length of each kind of microtubule decreased, with kinetochore microtubules (which are the longest, presumably because their plus ends are stabilized at kinetochores) experiencing the largest percentage decrease (approximately 30%; Figure 7B). The number of augmin-nucleated microtubules increased from zero to more than 1000 and the number of pole-nucleated microtubules decreased from around 500 to 400, while the number of kinetochore-nucleated microtubules remained approximately constant, as expected (Figure 7C). We also tested the effect of varying the augmin binding and nucleation rates as well as the diffusion coefficient on the spindle organization (Figure S6). Increasing binding and nucleation rates (Figure S6A–L) resulted in similar effects to increasing source rate. Increasing the diffusion coefficient (Figure S6M-R), however, had the opposite effect on spindle length. Indeed, the increased mobility, which occurs in random directions, is expected to diminish the chance of augmin to encounter and bind microtubules, effectively decreasing the amplification activity.

**Figure 7.**
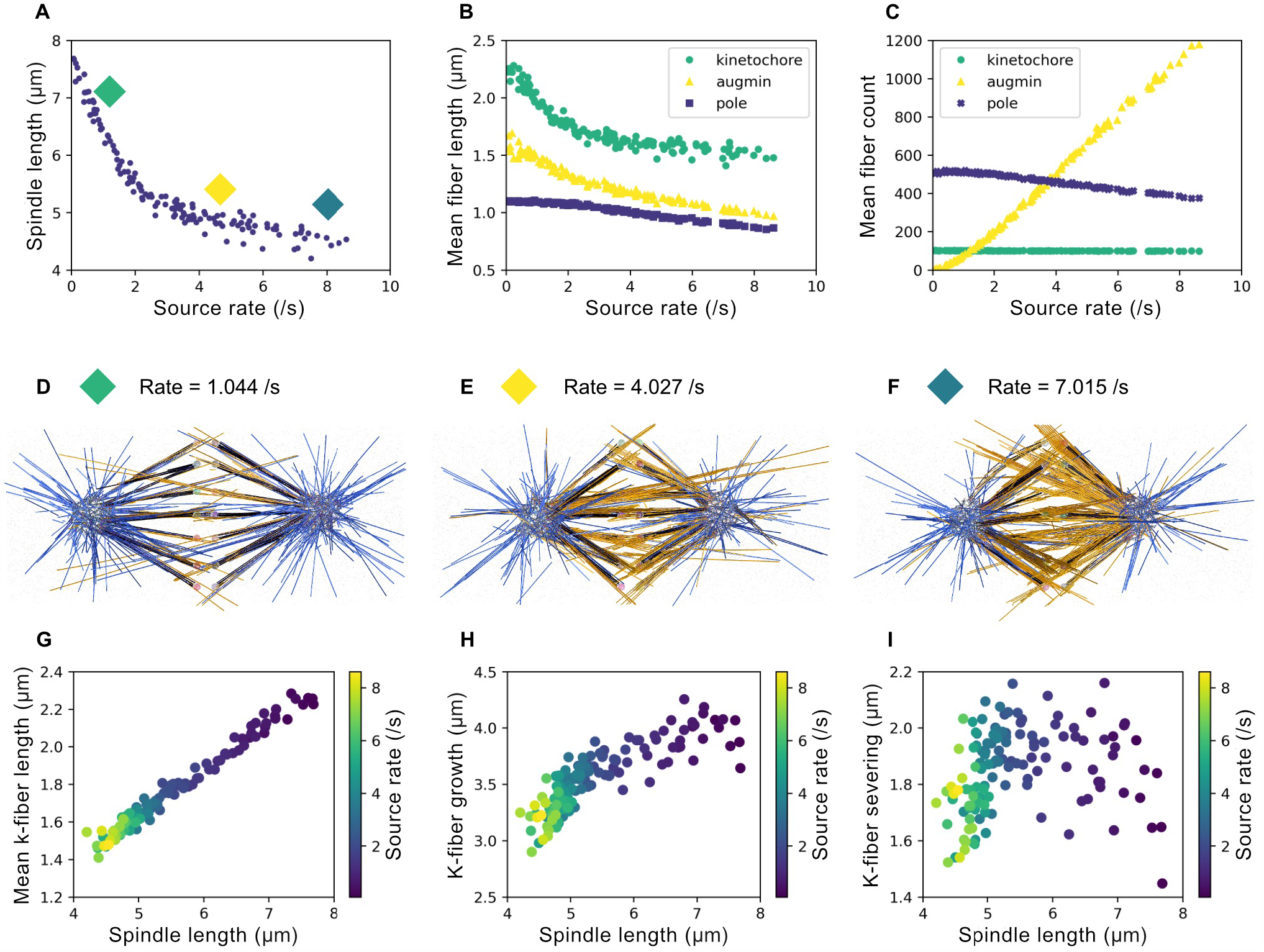
The amount of augmin controls spindle length and organization in the simulation. (A–C) Some key spindle properties as a function of the augmin source rate (/s). All temporal means are taken over the last half of the simulation, 500s < t < 1000s. (A) The mean spindle length (μm) decreases with augmin source rate (/s). The spindle length is measured as the distance between the center-of-masses of the left and right groups of condensates. The green, yellow, and blue diamonds indicate the three examples shown in (D). (B) Mean lengths (μm) of each group of fibers nucleated at kinetochores (green circles), by augmin (yellow triangles), and at poles (purple squares). (C) Mean number of fibers of each type. (D–F) Visualization of simulated spindles at the final time t = 1000s, for augmin source rates as indicated. Kinetochore microtubules are black, augmin-nucleated microtubules are orange, and pole-nucleated microtubules are blue. Kinetochores are variously-colored spheres near the metaphase plate. (G–I) Relationships between kinetochore-fiber (k-fiber) properties and the spindle length (μm), with data points colored according to the augmin source rate (/s). All quantities are means over the last half of the simulation, 1000s < t < 2000s, and k-fiber quantities are averaged over all k-fibers. (G) Mean k-fiber length (μm), (H) mean growth (μm) at k-fiber plus ends, and (I) mean severing (μm) at k-fiber minus ends.

## Discussion

In this work, we have combined computer simulations with experimental approaches to advance our knowledge of spindle formation in plants. We have identified CDKB1 in conjunction with CYCB3;1 as a major regulator of the Arabidopsis mitotic spindle. Until now, little was known in plants about how cell cycle regulators control spindle formation. Based on their role in microtubule organization^17^, we had initially expected that B1-type cyclins together with their CDK partners, mostly CDKB2s, would be involved in the regulation of spindle shape and organization. However, no obvious spindle defects were found in the most severe mutant combination *cycb1;1 cycb1;2*. Although we cannot rule out that other members of the B1 class participate in spindle architecture, CYCB1s seem mostly involved in other aspects of chromosome segregation, like connection of spindle microtubules to kinetochores^17^. Accordingly, the B1-type cyclin from humans binds to and supports the localization of a member of the spindle assembly checkpoint (SAC), MAD1, at the kinetochore^35^. With respect to plant B1-type cyclins, it will be interesting to explore whether they have a similar role in regulating kinetochore proteins and/or the SAC, especially given that the core SAC machinery appears to be functionally-conserved in Arabidopsis albeit in an adapted manner^36,37^.

CDK-cyclin complexes have been previously implicated in the direct control of spindle morphogenesis in other organisms. For instance, the Cdk1-cyclin B1 complex from humans is known to phosphorylate importin-α1 to inhibit its function, and release spindle assembly factors, such as TPX2, to promote spindle formation^38^. Furthermore, mutations in the budding yeast CDK1 (Cdc28) as well as simultaneous depletion of all budding yeast B-type cyclins also result in abnormal spindle assembly, which mirrors our findings with CYCB3;1 in Arabidopsis. More specifically, budding yeast cells impaired in Cdc28/B-cyclin function have duplicated spindle pole bodies (SPBs) that fail to separate^39^. The Cdc28/B-cyclin complex specifically phosphorylates yeast kinesins-5 Kip1 and Cin8, and this phosphorylation plays a role in promoting SPB separation and spindle assembly^40^. Although plants lack a discernable MTOC at the spindle stage like SPBs or centrosomes, here we found that γ-tubulin (a major component of SPBs and centrosomes) distribution is likewise impaired in *cycb3;1* and *cdkb1* mutants. In Arabidopsis, no less than 23 kinesins are expressed in mitosis, among which many have potential CDK phosphosites^41^. Whether B-type cyclins are involved in the phospho-control of such mitotic kinesins and help establish spindle bipolarity in plant cells remains to be seen.

### The role of CDKB1 and CYCB3;1 in spindle organization

Here, we found that the CYCB3;1-CDKB1 complex is involved in spindle morphogenesis, at least partly through phospho-regulation of the augmin complex member EDE1. Interestingly, tubulin turnover does not seem to be affected in the non-phosphorylatable version of EDE1 we analyzed; hence, we propose that the elongated spindle phenotype we observed is mostly due to an altered frequency and/or pattern of microtubule-dependent microtubule nucleation within the spindle. If tubulin availability in a cell limits spindle length, spindles can become longer when augmin function is affected because the amount of free tubulin increases, as does the polymerization speed of the remaining spindle microtubules. Indeed, in our simulations, spindles became shorter in response to increasing augmin source rate. As the augmin source rate increased from 0.065 /s to 8.6 /s, the amount of free tubulin (measured as the microtubule length equivalent) decreased from 3300 μm to 2500 μm, which means that the microtubule growth speed decreased by about a third relatively to its base speed. Consequently, kinetochore microtubules became shorter, contributing to shortening the spindle (Figure 7G–I). Furthermore, pole-nucleated microtubules were longer and more abundant with lower levels of augmin in our simulations, fitting our observation of a higher and more prominent number of astral microtubules in *cdkb1* mutants. Perhaps parallel nucleation and other augmin-independent nucleation pathways become more common in the mutants we studied, further contributing to the change in spindle shape we observed as previously suggested for the *ede1-1* mutant^8^. Additionally, since the augmin complex nucleates microtubules that generally preserve the polarity of their mother microtubules^6^, the astral microtubules in *cycb3;1* and *cdkb1* represent further evidence of a deficient augmin activity.

Why do *cycb3;1* and *cdkb1* mutants display spindles with an altered distribution of γ-tubulin, biased towards the poles? At the prophase stage, the pro-spindle is present normally as two polar caps rich in γ-tubulin at the nuclear envelope^42^, and this structure seems unperturbed in the analyzed mutants. Following NEB, augmin has been shown to critically bind to and amplify the number of microtubules to assist spindle formation^12^. Augmin likely translocates γ-tubulin from the spindle poles (which form from remnants of the polar caps following NEB) towards spindle microtubules in the midzone. In the *cycb3;1* and *cdkb1* mutants, however, a faulty augmin-mediated redistribution of γ-tubulin upon NEB likely results in the accumulation of γ-tubulin at the spindle poles.

Since the spindle defects seen in *cdkb1* double mutants are stronger than in *cycb3;1* mutants, it seems probable that CDKB1s operate with other cyclins to control spindle morphology. CDKB1s may also be involved in the establishment of the cortical division site, since we often observed cells without a PPB in *cdkb1* mutants (Figure 3E and 3F), although we did not examine this further.

### Robust sister chromatid separation by highly disorganized spindles

It is surprising that the spindles of *cdkb1* mutants, albeit highly disorganized, did not impair chromosome segregation, but rather affected the duration of spindle stages. It is likely that the action of the SAC ensured proper spindle-kinetochore attachments and bipartite sister chromatid segregation by delaying anaphase onset^36,43^.

A recent study in human spindles made use of a co-depletion of both the SAC factor Mad2 and HAUS6 to circumvent extensive mitotic divisions and study the effect of depleting augmin on chromosome segregation without the surveillance mechanism mediated by the SAC^44^. An interesting experiment for the future would be to combine mutations in SAC components with either the *cycb3;1* or *cdkb1* mutants to investigate how severely chromosome segregation is disrupted when both augmin and the SAC are impaired in plants.

### Basic molecular mechanisms guiding spindle organization

Here, we have modelled a spindle in three dimensions with increased realism in comparison to previous work and including new factors such as augmin and kinesin-14^19,45^. Whereas a quantitatively accurate model of the *Xenopus* spindle has not yet been achieved due to its size, the smaller size of the Arabidopsis spindle meant it was possible to simulate all of its microtubules within a reasonable computational time. Modelling an Arabidopsis mitotic spindle in particular was interesting because it has an intermediate size that is ideal for simulations when compared to smaller fission yeast or larger *Xenopus laevis* spindles and because plant spindles lack key molecular players seen in animals. For instance, there is only limited evidence of a NuMA homolog in plants^12,46^ and, hence, the pole organization in our simulation differs from the NuMA-organized spindle poles that have previously been employed^19,45^. Plants also lack the molecular motor dynein, which was also not included in our simulation, but possess an astonishing number of kinesins, including several expressed in mitosis, that likely take over some of dynein’s functions^47^.

With this work, we shed light on molecular mechanisms governing spindle organization in plants that are likely relevant for other eukaryotic groups as well. Our simulation will serve as a foundation for understanding spindle organization in other species, thus advancing our knowledge of how cells ensure a robustly-functioning spindle structure to separate their chromosomes in cell divisions and thereby proliferate.

## Acknowledgments

We thank Dr. Roland Thünauer for technical support at the Advanced Light and Fluorescence Microscopy facility at the CSSB (DESY, Hamburg), Anne Harzen at the MPI for Plant Breeding Research for LC-MS/MS analysis in the identification of EDE1 phosphosites and the VIB Proteomics Core for the LC-MS/MS analysis of the AP-MS samples. We thank the Cambridge Research Computing services, specifically for HPC used for this study. This work was supported by the HFSP grant RGP0023/2018 to A.S. and D.B.; F.N., H.S. and C.J. were supported by the Gatsby Charitable Foundation (Grant PTAG-024) and the European Research Council (ERC Synergy Grant, project 951430).

## Author contributions

M.R.M. and A.S. conceived the experiments. M.R.M., P.C. and L.B. performed experiments and statistical analyses. F.N., H.S. and C.J. constructed the spindle simulations and analyzed them. E.W. fixed and processed plant samples for TEM and performed TEM imaging. M.P., K.B. and D.B. performed the wholemount immunolocalization of α-tubulin and KNOLLE and corresponding statistical analyses. S.C.S. and H.N. performed the mass spectrometry experiment and data analysis of the *in vitro* kinase assays. E.V.D.S. and G.D.J. performed the AP-MS of CYCB3;1 and corresponding statistical analyses. M.R.M. and A.S. analyzed the data. M.R.M. and A.S wrote the manuscript.

## Figure legends

**Figure S1.**
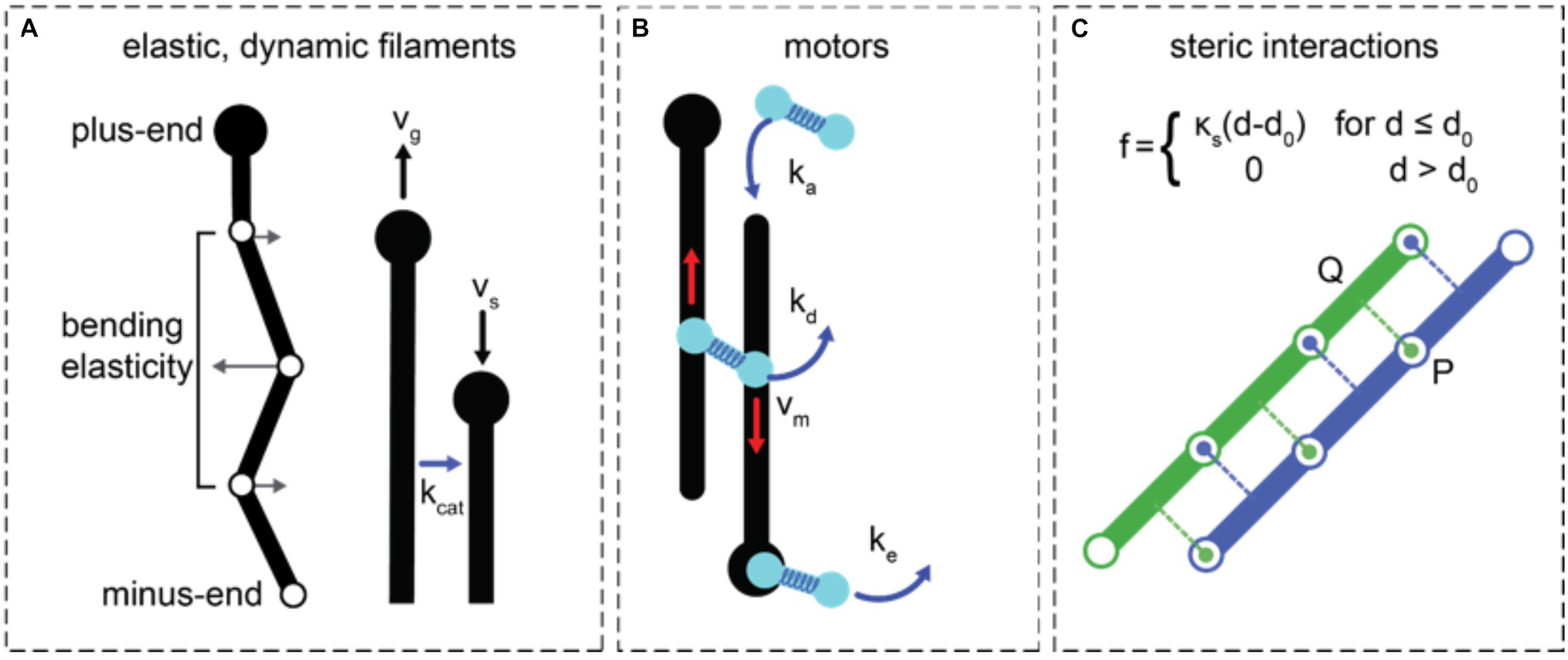
Essential elements of Cytosim. (A) Microtubules exhibit dynamic instability. They are discretized into points connected by inextensible segments such that microtubules can bend but do not stretch. Points are subjected to forces from bending elasticity, steric interactions, and crosslinking motors (if present). (B) Motors consist of two motor entities. Each motor entity can bind, unbind, and move along a microtubule. Crosslinking motors exert forces on the microtubules they connect via a Hookean spring-like link. (C) Steric interactions are calculated for each model point of a microtubule. For example, a line from P, a point on the blue microtubule, is projected onto the nearby segment of the green microtubule at Q. The line PQ is orthogonal to the green microtubule. An equal and opposite force is applied to the green and blue microtubule along PQ such that the steric forces acting on a pair of microtubules are symmetric and sum to zero.

**Figure S2.**
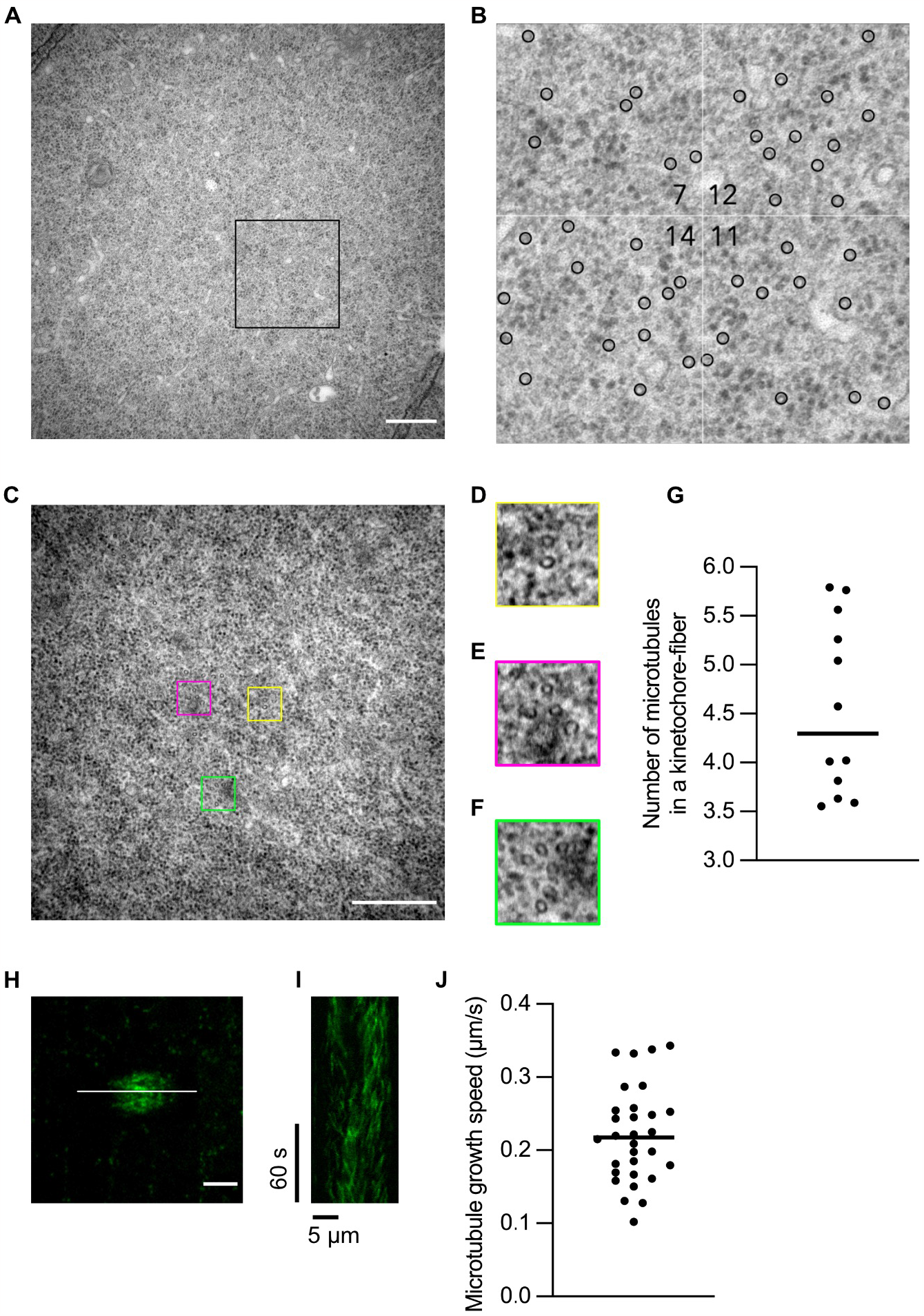
Experimentally-determined spindle parameters. (A) A cross section of a root spindle imaged by TEM. The area which was used to count microtubules is indicated with a box. Scale bar 0.5 μm. (B) Close-up corresponding to 1 μm^2^ indicated in (A). (C) A cross section of a root spindle imaged by TEM. Bundles with different microtubule numbers are indicated with colored boxes. Scale bar 0.5 μm. (D–F) Close-ups of microtubule bundles observed in (C). (D) A bundle of two microtubules. (E) A bundle of four microtubules. (F) A bundle of six microtubules. (G) Quantification of the number of microtubules in kinetochore-fibers measured from confocal microscopy images of root spindles stained against α-tubulin (n = 12 kinetochore-fibers from four different cells). The plotted line indicates the median. (H) A root spindle of a plant expressing *PRO*_*EB1b*_*:EB1b:GFP*. The line indicates the axis from which the kymograph in (I) was plotted. Scale bar 5 μm. (I) A kymograph generated from the line in (H). (J) Quantification of the microtubule growth speed from three independent spindles (n = 10 microtubules per spindle). The plotted line indicates the median.

**Figure S3.**
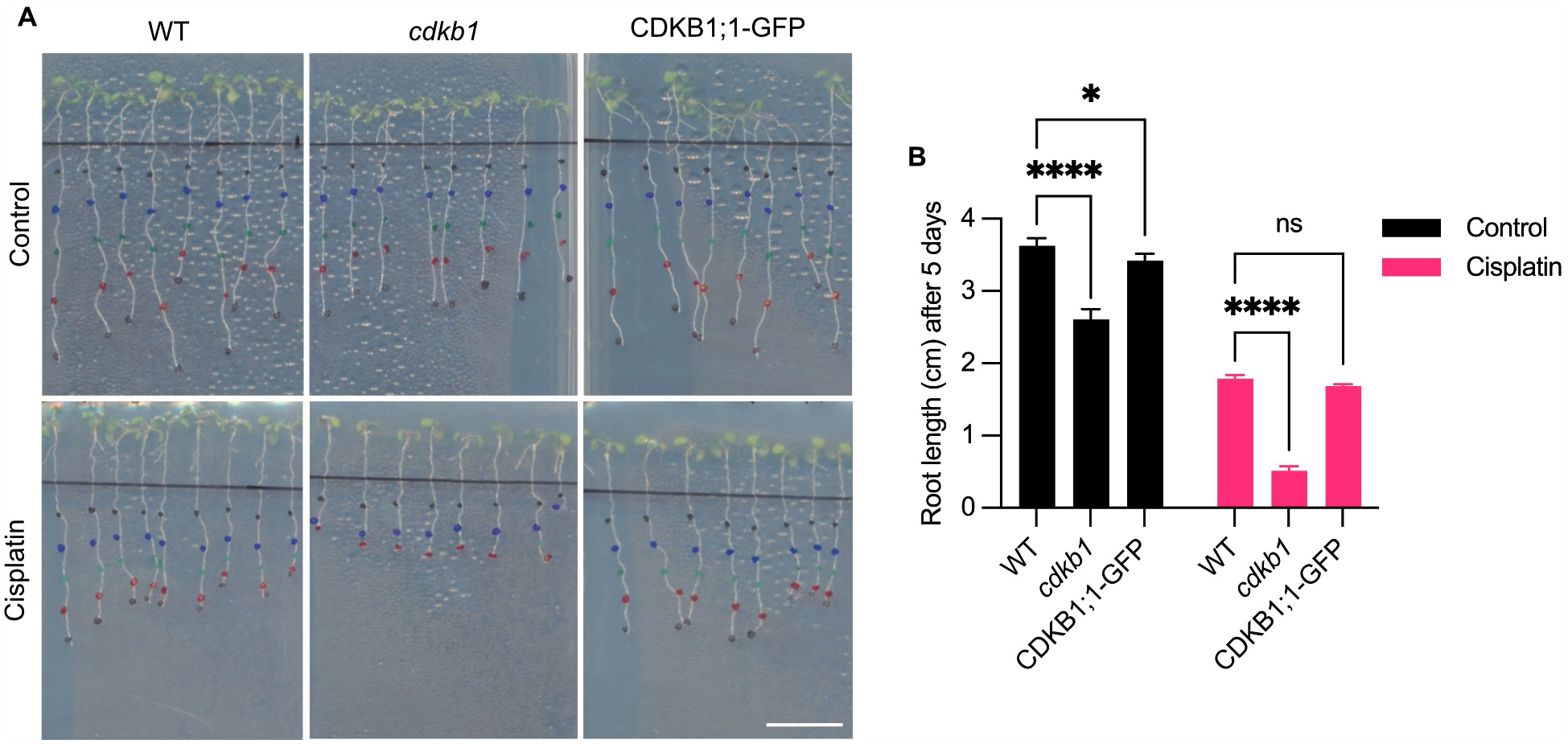
The CDKB1;1-GFP reporter largely rescues the root growth of *cdkb1* with and without cisplatin. (A) Pictures of seedlings of WT, *cdkb1* and *cdkb1* rescued by CDK1B1;1-GFP grown on ½ MS (control, top) or cisplatin (bottom) for five days. Scale bar 1 cm. (B) Quantification of root growth of WT, *cdkb1* and *cdkb1* rescued by CDKB1;1-GFP grown on ½ MS (control) or cisplatin for five days. Three replicates were performed with 10 plants per genotype per condition. Graph indicates mean ± SD of the three replicate average values. The level of significance was determined by a two-way ANOVA followed by Tukey’s multiple comparisons test (* P < 0.05 and **** P < 0.0001; ns depicts a non-significant difference).

**Figure S4.**
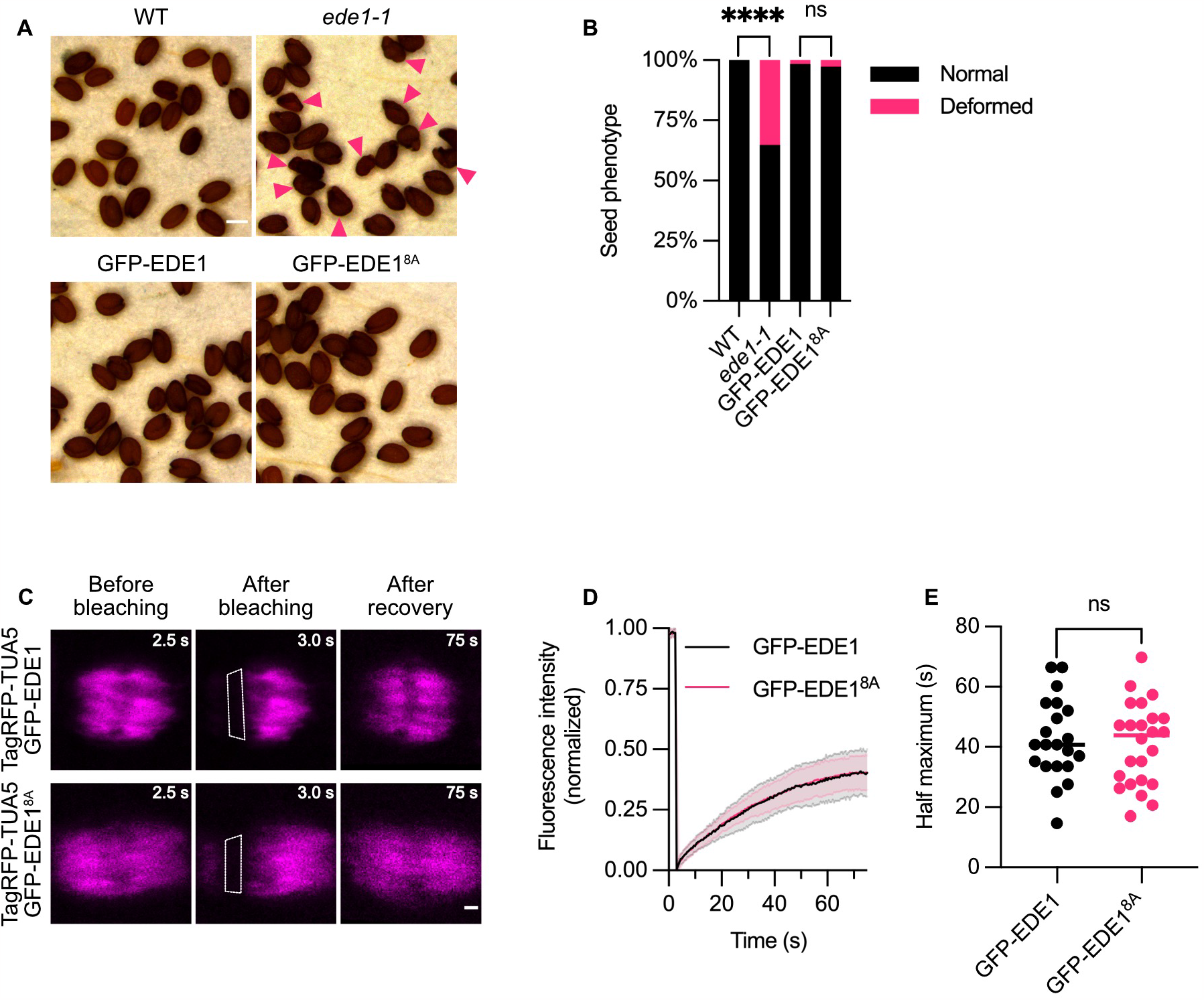
The GFP-EDE1^8A^ construct fully rescues the seed abortion of the *ede1-1* mutant and does not significantly affect the microtubule dynamic instability at the spindle. (A) Pictures of the seeds from WT and *ede1-1* as well as *ede1-1* mutants rescued by GFP-EDE1 and GFP-EDE1^8A^. Scale bar 200 μm. (B) Quantification of the seed abortion of the seeds depicted in (A) in WT (n = 158) and *ede1-1* (n = 190) as well as *ede1-1* mutants rescued by GFP-EDE1 (n = 189) and GFP-EDE1^8A^ (n = 188). (C) Confocal laser-scanning micrographs of root cells in which the FRAP assay of spindles tagged by TagRFP-TUA5 in the *ede1-1* mutant rescued by GFP-EDE1 or GFP-EDE1^8A^ was performed. The white dashed box represents the area that was bleached by the laser. The time is indicated on the top right of the images in seconds. Scale bar 1 μm. (D) Quantification of the fluorescence intensity recovery over time following bleaching of spindles in *ede1-1* mutant root cells tagged by TagRFP-TUA5 and rescued by GFP-EDE1 (n = 21) or GFP-EDE1^8A^ (n = 24). (E) Quantification of the half maximum values in seconds of fluorescence recovery in *ede1-1* mutants rescued by GFP-EDE1 (n = 21) or GFP-EDE1^8A^ (n = 24). The median values were plotted as a line for each genotype. Average half maximum of 42.52 s ± 13.32 for GFP-EDE1 and 40.90 s ± 13.71 for GFP-EDE1^8A^. Comparison on graph: *P* = 0.6903. The level of significance was determined by a two-proportion z-test followed by Bonferroni correction in (B) and an unpaired t test in (E) (**** P < 0.0001; ns depicts a non-significant difference).

**Figure S5.**
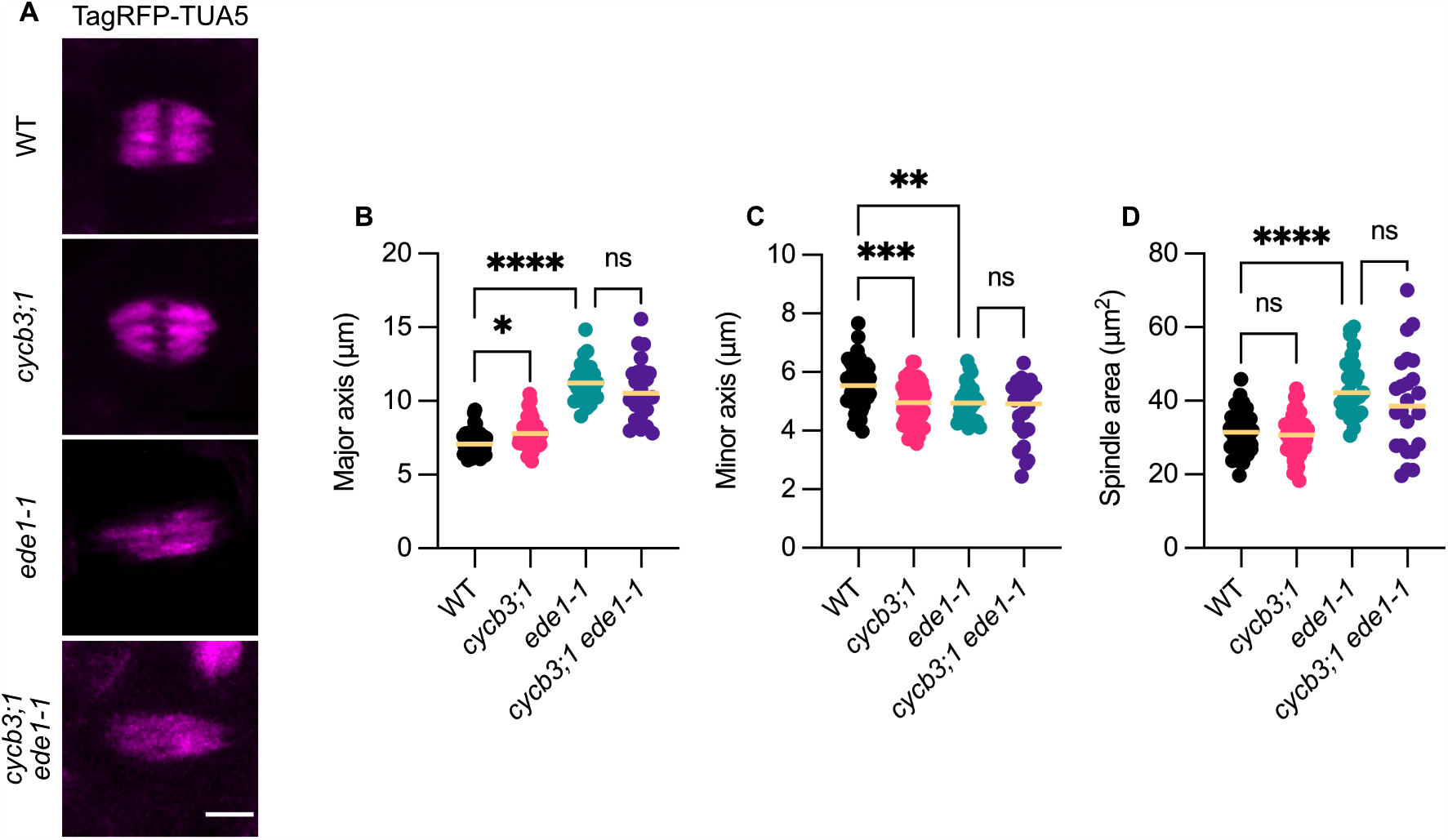
A double mutation in both *CYCB3;1* and *EDE1* does not further increase the spindle phenotype in comparison to the *ede-1* mutant. (A) Confocal laser-scanning micrographs of TagRFP-TUA5-tagged microtubules in root cells at the spindle stage of WT, *cycb3;1, ede1-1* and *cycb3;1 ede1-1* plants. Scale bar 5 μm. (B–D) Quantification of the spindle major axis (B), minor axis (C) and area (D) in root cells of WT (n = 50), *cycb3;1* (n = 51), *ede1-1* (n = 27) and *cycb3;1 ede1-1* (n = 24) plants. Median values were plotted as a line for each genotype. The level of significance was determined by an ordinary one-way ANOVA followed by Šídák’s multiple comparisons test (* P < 0.05, ** P < 0.01, *** P < 0.001, **** P < 0.0001).

**Figure S6.**
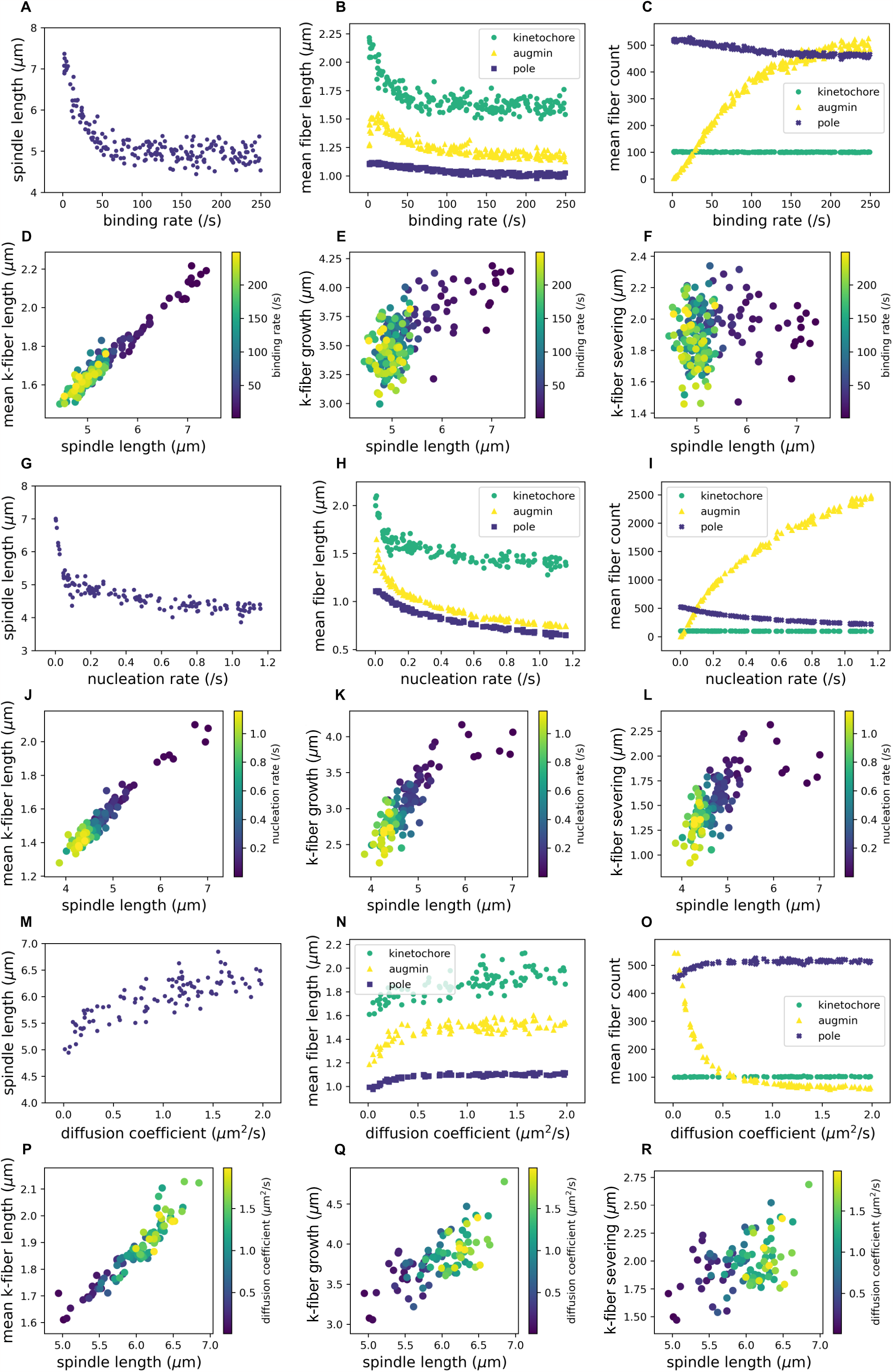
Varying other augmin parameters leads, in general, to a similar trend in spindle organization to varying augmin source rate. (A–C) Relationship between spindle properties and the augmin binding rate (/s). All temporal means are taken over the last half of the simulation, 500s < t < 1000s. (A) Mean spindle length (μm) against augmin binding rate (/s). The spindle length is measured as the distance between the center-of-masses of the left and right groups of condensates. (B) Mean lengths (μm) of each group of fibers nucleated at kinetochores (green circles), by augmin (yellow triangles), and at poles (purple squares). (C) Mean number of fibers of each type. (D–F) Relationships between kinetochore-fiber (k-fiber) properties and the spindle length (μm), with data points colored according to the augmin binding rate (/s). All quantities are means over the last half of the simulation, 500s < t < 1000s, and k-fiber quantities are averaged over all k-fibers. (D) Mean k-fiber length (μm), (E) mean growth (μm) at k-fiber plus ends, and (F) mean severing (μm) at k-fiber minus ends. (G–I) Relationship between spindle properties and the augmin nucleation rate (/s). All temporal means are taken over the last half of the simulation, 500s < t < 1000s. (G) Mean spindle length (μm) against augmin nucleation rate (/s). The spindle length is measured as the distance between the center-of-masses of the left and right groups of condensates. (H) Mean lengths (μm) of each group of fibers nucleated at kinetochores (green circles), by augmin (yellow triangles), and at poles (purple squares). (I) Mean number of fibers of each type. (J–L) Relationships between kinetochore-fiber (k-fiber) properties and the spindle length (μm), with data points colored according to the augmin nucleation rate (/s). All quantities are means over the last half of the simulation, 500s < t < 1000s, and k-fiber quantities are averaged over all k-fibers. (J) Mean k-fiber length (μm), (K) mean growth (μm) at k-fiber plus ends, and (L) mean severing (μm) at k-fiber minus ends. (M–O) Relationship between spindle properties and the augmin diffusion coefficient (μm^2^/s). All temporal means are taken over the last half of the simulation, 500s < t < 1000s. (M) Mean spindle length (μm) against augmin diffusion coefficient (μm^2^/s). The spindle length is measured as the distance between the center-of-masses of the left and right groups of condensates. (N) Mean lengths (μm) of each group of fibers nucleated at kinetochores (green circles), by augmin (yellow triangles), and at poles (purple squares). (O) Mean number of fibers of each type. (P–R) Relationships between kinetochore-fiber (k-fiber) properties and the spindle length (μm), with data points colored according to the augmin diffusion coefficient (μm^2^/s). All quantities are means over the last half of the simulation, 500s < t < 1000s, and k-fiber quantities are averaged over all k-fibers. (P) Mean k-fiber length (μm), (Q) mean growth (μm) at k-fiber plus ends, and (R) mean severing (μm) at k-fiber minus ends.

## Material and Methods

### Arabidopsis root mitotic spindle simulation

Mitotic spindles were simulated using Cytosim, an Open-Source project (gitlab.com/f-nedelec/cytosim). Here, we provide an overview of our methods which are based on Brownian dynamics. The numerical aspects (integration, stability) were described previously^18^. Further to this publication, accessibility of the source code should enable the full analysis of our methods, and reproducibility of the results. In brief, microtubules are modeled as incompressible bendable filaments having the persistence length of microtubules, in a medium characterized by a viscosity as measured for cells^48^. Microtubules are represented by vertices distributed regularly along their length. Connections between microtubules, and forces such as steric interactions are represented by Hookean links between the filament’s vertices. The forces are linearly interpolated to adjacent vertices on the filament when a link is formed between two vertices. The evolution of the entire network is simulated by solving the equation of motion for successive small-time intervals, updating this equation as the motors move to different positions on the filaments, and motor and crosslinkers bind or unbind, and microtubules grow, shrink, vanish, or are created. In essence, the movement is defined by an over-damped Langevin equation: 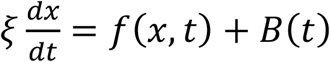, for a large multivariate vector *x*, where the right-hand terms are elastic and random forces respectively, and *ξ* is a diagonal matrix of drag coefficients calculated using Stokes’ law from the viscosity of the fluid and the dimensions of the objects. Such an equation accurately describes the motion of micrometer-sized objects in a fluid that is dominated by elastic and viscous forces. In addition to Brownian motion at each positional coordinate, the equation includes the bending elasticity of the filaments and the elastic terms associated with the molecules forming bridges between two filaments. The differential equation involving all the coordinates of all vertices is solved using a first-order semi-implicit numerical integration scheme that is numerically very stable. Moreover, at each time step, a variety of biochemical processes are modelled as first-order stochastic processes: activation, binding and unbinding, nucleation, microtubule dynamic instability.

The cell volume is fixed and cylindrical, with a length 11μm and diameter 5μm, symmetric around the x-axis. The edges of the cell induce microtubule plus ends to stall. With this assumption, no confinement forces were needed. Microtubules thus do not track the cell edges.

Microtubule Nucleation. Microtubules are nucleated by three pathways:

N1. Kinetochores

N2. Pole-induced

N3. Augmin-mediated

Each pathway is constituted of a fixed number of nucleators entities, with properties adjusted according to the pathways that is represented: kinetochore and pole associated nucleators are anchored to beads, while the augmin-mediated nucleator is part of a diffusible complex. Each nucleator may only nucleate one microtubule at a time, and would remain inactive until this microtubule vanishes, or the nucleator detaches from it.

For pathway N1 (kinetochores), each kinetochore harbors 5 nucleators, and their nucleation rate is fixed and unregulated. Moreover, the kinetochore-based nucleator will remain attached at the plus end of the microtubules, while for the other pathways the nucleator remains attached to the minus end.

Nucleation pathway N2 (poles) consists of nucleators attached to the beads that form the condensate at the spindle pole (see below).

For nucleation pathway N3 (augmin-mediated), individual augmin entities are generated on a random position on the surface of the kinetochores with a fixed source rate. These augmin entities have a finite lifetime characterized by a constant molecular rate. This is implemented using a timer for each augmin entity, initialized with t = −*log*(*θ*^+^)/*R*, where R=5/s is the deactivation rate and *θ*^+^ a random number in [0,1]. Augmin entities are deleted if their timers reach zero. During its lifetime, an augmin entity diffuses freely, and may bind to existing microtubules within its binding range, with the prescribed binding rate. A bound augmin stays fixed relative to the microtubule on which it is attached, until it unbinds. An augmin entity that is bound to a microtubule (the mother) will nucleate a new microtubule (the daughter) as determined by its nucleation rate. Unbound augmin do not nucleate. A daughter microtubule is orientated parallel to the mother microtubule, in the same direction. During the time that it is bound, the augmin entity is protected from deactivation (the internal timer is frozen). The timer restarts if the augmin detaches from the microtubules to which it is docked. These assumptions are intended to capture the control of the augmin activity by the Ran pathway^49^, where the RanGTP complex is generated at the surface of the chromatin by RCC1 and deactivated elsewhere by RanGAP, forming a sharp gradient of active Ran around the chromosomes^50^. Our assumptions capture the essential condition that daughter microtubules are nucleated parallel to their mother microtubule^4^, in the vicinity of the chromosomes^51^, and that augmin can be transported by fluxing microtubules^52^.

We used a single scalar parameter (noted as *γ*) to model the fact that pathways 2 and 3 share the same molecular nucleator gamma-tubulin. When a nucleator from these two pathways is active, its nucleation rate is multiplied by (1 − *Nγ*), where N is the number of microtubules in the system. In this way nucleation is reduced as microtubules become more numerous until it vanishes for *N* = 1/*γ*. We used *γ* = 0.0001, corresponding to a maximum of 10000 microtubules, which is much above the actual number of microtubules in the simulation (∼2000), and this limit is not reached. However, this assumption connects the nucleation activities of pathways 2 and 3, with the effect of reducing the number of pole-nucleated microtubules if the augmin activity is increased.

Microtubules are nucleated with an initial length *L*_0_ = 32*nm* with their plus ends in the growing state, and undergo dynamic instability at the plus ends. The minus ends are static. Dynamic instability at the plus end is implemented following a stochastic model of the GTP cap that protects microtubules from catastrophes^53^. The instantaneous microtubule growth speed is set dynamically from the total length of the microtubules at a given time point i.e. 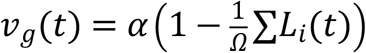 where *α* is the maximum growth speed, ∑*L*_*i*_(*t*) is the total length of all microtubules at time *t* and the constant *Ω* represents the total available tubulin pool, expressed in MT length (4000μm). These assumptions intend to represent conditions in which the amount of tubulin from which microtubules polymerize is finite. The growth speed of individual microtubule is further reduced in the presence of an antagonistic force, *f*_*a*_ < 0, by an exponential factor, 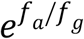, where *f*_*g*_ > 0 is a characteristic “growing force” parameter^54^. This factor is always applied, but we believe that it is insignificant for the simulations presented in this work, because cell-edge induced forces were not enabled. We instead assumed that microtubules would stall upon contacting the cell edge, only growing at a fraction of their speed in the cytoplasm; specifically, the growth speed is divided by 10. With this assumption, we recover the conditions in plant cells, where microtubules are not observed to track the edges of the cell, but instead Eb1 comets vanish as they reach the cell edge. Given that the stochastic model of dynamic instability is very dependent on the rate of tubulin addition, microtubules contacting the cell edge thus rapidly undergo catastrophes in the simulation, as observed *in vivo*. Microtubules shrink at a constant shrinkage speed *v*_*s*_ and do not undergo rescues. Any microtubule shorter than 24 nm is deleted. After a shrinking microtubule has vanished, its nucleator is free to nucleate again.

Microtubules experience steric interactions. They repel each other via a soft-core interaction that is repulsive with a diameter *d*_0_ = 50*nm*:

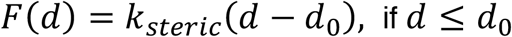

where *d* is the distance between the two interacting filaments. This force is applied at every filament vertex that is within the steric diameter of another filament segment. It acts primarily in the direction orthogonal to the filament axis and will not prevent filaments from sliding along each other. Steric forces interfere in this way minimally with the movements induced by crosslinking motors such as Kinesin 5 (Figure S1) but will induce parallel microtubules to separate their center lines *d*_0_ = 50*nm* apart.

Moreover, a weak force is added to bring the microtubules closer to the x-axis (parameter ‘squeeze’). This force promotes the formation of the spindle poles by focusing the kinetochores fibers on the x-axis. The force magnitude is implemented as *f*(*u*) = *F*_*ε*_ *tanh*(*u*/*R*_*z*_), with 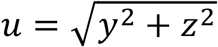 and *F*_*ε*_ = 0.05*pN* the maximum magnitude of the force, and *R*_*z*_ = 3*µm* is the range at which it plateaus. The force is applied only at the minus ends, to all microtubules. This force is directed towards the x-axis, with no component parallel to the x-axis: 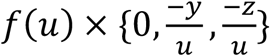.

Kinetochores are represented by spherical particles with a radius of 100 nm. The 20 kinetochores associated with the 10 chromosomes are placed such as to form a regular metaphase plate in the middle of the cell. Ten kinetochores are placed a *x*= 0.25*µm*, while the other ten are placed at *x*= −0.25*µm*, in a mirror configuration (same *y* and *z* coordinates). The two sets of 10 kinetochores are distributed in the YZ plane such as to approximate a disc of uniform density. Specifically, 8 kinetochores are placed at the summit of a regular octagon with *y*^2^ + *z*^2^ = 2*µm*, and two kinetochores are placed inside this octagon at *y* = 0.65*µm* and *z* = 0.3*µm* and the symmetric position {−*y*, −*z*}. Each kinetochore is immobilized in translation with a Hookean link of stiffness 1000 pN/μm but is free to rotate. Thus, the metaphase alignment of the chromosomes is assumed in our model. Each kinetochore harbors 5 nucleation entities. Microtubules are allowed to grow from the kinetochores in the initialization sequence of the simulation, in the direction of the closest spindle pole (*e*.*g*., toward *x*> 0 for microtubules originating from kinetochores placed at *x*= 0.25*µm*). This favors the biorientation of all kinetochores in the initial configuration. The alignment of chromosomes in the metaphase plate, and the biorientation of kinetochores are two important aspects of mitotic spindle assembly that were intentionally left aside for future work, to focus on the question of how the length of the spindle is regulated by augmin.

Each kinetochore has 5 nucleating entities (ndc80) located on a cap directed towards the closest pole. Each entity may nucleate one microtubule and remains attached to its plus end until spontaneous detachment occurs, which is set at a rate of 0.01s^-1^. The nucleation rate of 1s^-1^ implies that kinetochores have 5 microtubules attached to them most of the time. If the kinetochore unbinds, the associated microtubule plus end is set in a shrinking state and will thus rapidly vanish since there is no rescue. Kinetochores regulate the plus end dynamics of microtubules to which they are attached. The minus ends are not affected. A kinetochore-attached microtubule plus end grows slower than that of a regular microtubule, and its growth speed is regulated by force on the plus end *f* (the force in the ndc80 entity). Specifically, 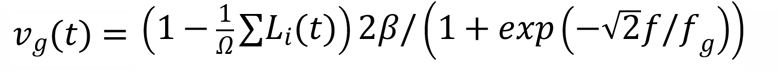, where *β* = *α*/5 (the amplitude of the reduction, 5, is set by the parameter ‘stabilize’) and where *f*_*g*_ > 0 is the microtubule’s characteristic “growing force”. Compared to other microtubules, the kinetochore suppresses catastrophes, reduces average growth by a factor 6, and regulates growth upon force with the factor 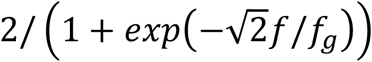, which by construction is in [0,2]. Pulling forces will accelerate microtubule growth up to a factor 2, while pushing forces will reduce growth by a significant fraction, if the force reaches *f*_*g*_.

Each spherical particle used to represent a kinetochore contains three vertices on its surface, constituting, together with the center point, a local reference frame that provides orientation in space. The ndc80 entities are placed with respect to this local reference frame, such as to form a small cluster (a ‘polar cap’) on one side. This cap is initially orientated towards the closest spindle pole. Kinetochores and associated microtubules are linked by Hookean links. A first type of link constrains the position of the plus end to match the position of the “ndc80” entity on the surface of the kinetochore. This link is of zero resting length and stiffness 222 pN/μm. A second type of link (parameter ‘anchor_stiffness’ 44 pN/μm) is used to align all the microtubules from one kinetochore, in the direction of its cap. This link is formed between the vertex of the microtubule, that is just before the plus end, and a matching virtual point built on the kinetochore reference frame, away from the kinetochore surface, at a distance equal to the separation of the microtubule vertex and its plus end. This way a geometrically simple but realistic configuration of microtubule attachment with the kinetochore is built.

Molecular motors. Kinesin-5 and Kinesin-14 are modeled as 2 linked units, forming a complex which can thus be unbound, attached to one microtubule, or attached to two microtubules. Complexes diffuse in the unbound state, can bind to one or two filaments and, when bound to two filaments, are modeled as Hookean springs with a resting length of 50 nm and various stiffness values as specified in the parameter table. Binding is determined by a rate within a binding range, and these two parameters are set following typical values for such molecules, initially measured for conventional kinesin. Subunits bind and unbind independently from each other but cannot bind to the same position on the same filament when they belong to the same complex. Diffusion of unbound motors is not modelled explicitly; it is assumed to be sufficiently fast that a uniform spatial distribution of unbound motors is maintained. The simulation only keeps track of the number of unbound motors, but not their positions and evaluates the average number of binding events per time step using the current total length of microtubules and the cell volume. This estimate is discretized using a Poisson distribution and the corresponding number of binding events is directly implemented by picking random positions along microtubules with uniform sampling (option ‘fast_diffusion’).

Molecular motor units. Whereas their binding and unbinding are discrete stochastic events, bound kinesins move deterministically on microtubules at a speed which is linearly proportional to load, given by *v* = *v*_*m*_(1 + *f*_*load*_ · *d*/*f*_*stall*_), where *d* is a unit vector parallel to the microtubule (in the direction preferred by the motor), *f*_*load*_ the force vector, *f*_*stall*_ > 0 is a characteristic stall force and *v*_*m*_ is the unloaded speed of the motor (positive for kinesin5 and negative for kinesin14). Note that with our conventions, forces that antagonize the motor preferred motion are directed opposite to *d*, hence a plus-end-directed motor is slowed down by forces directed towards the minus end. For a minus-end-directed motor, the unit vector *d* points toward the minus end. Motors detach from the microtubule side at a rate *k*_*off*_ and immediately from the microtubule ends. The detachment rates of motors are increased exponentially by the load on the motor and a characteristic unbinding force *f*_*unbind*_, according to Kramer’s law;*k* = *k*_*off*_*exp* (‖*f*_*load*_‖/*f*_*unbind*_).

Kinesin-5 is modelled as a pair of identical motor units connected by a Hookean spring-like link with resting length *d*_*m*_ and stiffness *K*_*m*_. This link can rotate freely at both attachment points, such that the angle between two crosslinked microtubules is unconstrained. If one motor of a pair is bound to a microtubule the other can bind to any microtubule within a range *r*_*b*_ at rate *k*_*on*_. To simulate the observed difference in Kinesin-5 affinity to parallel vs. antiparallel microtubules configurations^55^, we used two separate kinesin-5 entities: a ‘antikin’ that may only bind antiparallel configurations and a ‘parakin’ that may bind to all the other configurations. The criteria defining parallel vs. antiparallel is based on the cosine of the angle formed between the direction vectors of the relevant microtubule segments (the dot product of the unit direction vector of the microtubules). The antiparallel motor may bind only if *cos*(*θ*) < −0.5, and the other motor if *cos*(*θ*) > −0.5. To simulate the observed differences, the ‘parakin’ as an unbinding rate of 0.1 s^-1^, whereas the ‘antikin’ has a lower unbinding rate of 0.025 s^-1^. The other characteristics of the two kinesin-5 subtypes are identical.

Kinesin-14 is composed of a minus-end-directed motor domain linked to a diffusible domain via a Hookean link. The minus-end-directed motors is modelled similarly to the plus-end-directed motor domains of Kinesin-5, with respect to load and detachment. The non-motor domain of Kinesin-14 may diffuse passively or be dragged along the side of a microtubule. It is characterized by a linear mobility coefficient *µ*. A domain that is under a force *f* transmitted through the Hookean link will move along the microtubule in the direction of the force with an average speed *µf*. In addition, it undergoes diffusion with a 1D diffusion constant *D*_1_ = *µk*_*B*_*T*, where *T* is the absolute temperature and *k*_*B*_ Boltzman’s constant (*k*_*B*_*T* = 4.2*nm. pN*). The movement in a time interval *τ* was implemented as 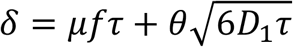, where *θ* is a random number uniformly distributed over [−1,1]. In contrast to the motor domain, the diffusible domain does not unbind immediately upon reaching the microtubule minus end. Instead, it keeps the same unbinding rate at the minus end than when located on the side of microtubules. Unbinding rates are however still modulated exponentially by the load according to Kramer’s law;*k* = *k*_*off*_*exp* (‖*f*_*load*_‖/*f*_*unfind*_). Given that it is linked to a slow minus-end-directed motor, a diffusible domain is unlikely to ever reach a growing plus end, but we have assumed anyhow that it would detach immediately at the plus end.

#### Spindle poles

The poles of the spindle in the simulation are made with discrete particles. Initially, 1000 particles are placed at *x*= 2*µm*, and another 1000 at *x*= −2*µm*. Two forces hold particles together and provide them with the ability to form a fluid phase within the cytoplasm: a specific pressure associated to the density of particles, and a surface tension. The pressure terms ensure that the beads remain separated by a distance roughly corresponding to maximal sphere packing density. The surface tension promotes the fusion of two droplets of beads that would come into contact, in our case leading to the collapse of the spindle into a monopole. The beads behave as a fluid phase and form compact droplets at the pole which remain mostly spherical and moves very little during the simulation. The density of the condensate appears uniform and close to the density value set as parameter (equal to *V*_*max*_).

The bead fluid subsystem is modelled following the ‘Smoothed Particle Hydrodynamics’ (SPH) method^56^. The SPH method, which was originally developed for astrophysics, integrates well with Cytosim after adaptation to the microscopic physics in which inertia is negligible. All particles are spherical with the same radius *R* = 64*nm*. We assumed a uniform mass density for the particles that is equal to that of the cytoplasm, such that we simply used the volume of each particle (*m*_*a*_ = 4*πR*^3^/3) and not their mass to weigh their contribution in the SPH sums. We note *h* the smoothing length scale (*h* = 303*nm*) and only used kernels with finite support, vanishing for distances d above *h*. The local density *ρ*_*a*_ is calculated using the standard 6^th^-order polynomial kernel *W*_*poly*6_(*d*) = *W*_6_ [*h*^2^− *d*^2^]^3^, where *W*_6_ = 315/64*πh*^9^ provides the normalization. With our simplification of density=1, the mass density estimated at particles is effectively a dimensionless volume fraction. A value of pressure for each bead is then calculated as *P*_*a*_ = *K*_*SPH*_. *max*(0, *ρ*_*a*_ − *V*_*max*_), where *V*_*max*_ is the desired density, set to the maximum volume fraction achieved for packed spheres 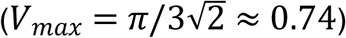, and *K*_*SPH*_ can be seen as a compressibility factor, a stiffness associated with the pressure. A pressure force is derived from the gradient of density, using Desbrun’s spiky kernel *W*_*spiky*_(*d*) = *W*_*S*_[*h* − *d*]^3^, where *W*_*S*_ = 15/*πh*^6^ ^57^, using Monagan’s symmetric formula (Eq. 3.3 in J. J. Monaghan. Smoothed particle hydrodynamics, 1992):

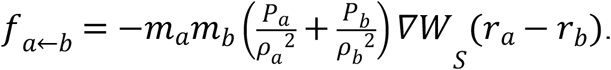

We used a cohesion kernel to model the surface tension^58^, with however a modified kernel:

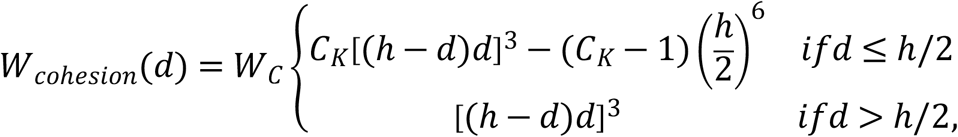

where W_C_ = 32/πh^9^ is for normalization. This kernel is continuous at *d* = *h*/2, and *C*_*K*_ = 275/19is adjusted to ensure that the force experienced by a particle located on the surface of a droplet of constant density would vanish, namely:

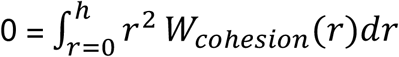

The cohesion force is calculated using a symmetric formula:

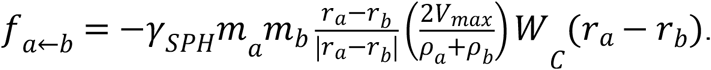

Using symmetric SPH formula ensures that the force will always balance, which is essential. The forces calculated per particle are then scaled by the drag coefficient of each particle (Stokes’ law: *ζ* = [6*πR*]^-1^) to obtain their instantaneous speed, from which a displacement is calculated. We use an explicit integration if the bead is unconnected with the microtubule system (*δx*= *ζfτ*). Otherwise, for instance if one of the bead’s binder is attached to a microtubule, the SPH force is added to Cytosim’s force engine as an explicit force term, such as to combine the SPH-calculated forces with the elastic forces associated with the links to microtubules. In any case, a random force is added to model the unbiased diffusion of beads, calibrated from their size (*D* = *ζk*_*B*_*T*), the viscosity and the time-step. The full details of our SPH implementation will be given in a separate article.

Three activities are associated with the beads forming the condensate at the spindle pole: microtubule binding, nucleation and severing. The microtubule binding activity is implemented by attaching discrete binding entities to the center of the beads forming the condensate. In the spindle simulations, each bead has ∼4 binders. These binders may only bind to microtubules near their minus ends, specifically at a location of the microtubule that is less than 250 nm away from the minus end, provided the distance to the bead center (where the binder is anchored) is lower than 64 nm. The nucleation activity is implemented by attaching one nucleator per bead (see above for the detailed description of the nucleation model). The microtubule severing activity is implemented similarly to the augmin complex: katanin entities are generated at the surface of the beads with a ‘source’ rate and destroyed stochastically with a constant rate of 8 s^-1^. Katanin entities are free to diffuse and to bind to microtubules during their lifetime. In this way a permanent gradient of severing activity is generated within and around the condensate. A Katanin entity is a complex made of two severing units. Each severing unit can cut a microtubule to which it is bound with a rate of 0.2s^-1^. Upon cutting the severing unit unbinds. The new plus end is created in the shrinking state, as widely observed^59^. The new minus end is static.

### Experimental determination of spindle parameters

For the estimation of the number of spindle microtubules, we analyzed TEM images of cross-sections of roots. We measured the number of microtubules in a 1 μm^2^ square. Next, we extrapolated this value to an area of approximately 16 μm^2^ for one half of the spindle. With this, we reached a value that varied between 576 and 1,408 microtubules for a full root spindle (n = 5).

For estimating the number of microtubules in kinetochore fibers, we counted the number of microtubules in bundles from TEM images and measured the fluorescence intensity of kinetochore fibers from spindles stained against α-tubulin compared to single microtubules in the same cell. For the fluorescence measurements, we drew a line across a single microtubule in Fiji and measured the integrated density divided by the area analyzed. Next, we measured the integrated density divided by the area analyzed in kinetochore fibers and divided that by the value obtained for a single microtubule to obtain an estimate of number of microtubules. This fluorescence intensity estimate was obtained from four different cells. We obtained similar values in both experimental approaches.

For determining the growth speed of microtubules, we generated kymographs using the KymographBuilder Fiji plugin (https://imagej.net/plugins/kymograph-builder) from roots of plants expressing an *EB1b*_*PRO*_*::EB1b-GFP* reporter that were imaged with a spinning disk microscope with a 0.5 s frame rate. Values were obtained for ten microtubules per spindle from three spindles, each from an independent plant.

### TEM of Arabidopsis root cross-sections

Roots were fixed with 2% glutaraldehyde in cacodylate buffer (75 mM, pH 7.0) for 3.5 h, postfixed with 1% osmium tetroxide at 4°C overnight. Samples were dehydrated through a series of graded acetone concentrations, 30% to 100%, and finally embedded in plastic according to Spurr^60^. Ultrathin sections were obtained with a ultramicrotome (Ultracut E, Leica-Reichert-Jung, Nußloch, Germany) and stained with uranyl acetate followed by lead citrate^61^. Sections were viewed with a LEO 906 E TEM (LEO, Oberkochen, Germany) equipped with the Wide-angle-2K (4Mpx.) Dual Speed CCD Camera (TRS, Moorenweis, Germany) using the software ImageSP-Professional to acquire, visualize, analyse, and process image data.

### Plant growth conditions

*Arabidopsis thaliana* seeds were grown on ½ MS medium (basal salt mixture, Duchefa Biochemie) containing 0.5% sucrose and 0.8% agar (plant agar, Duchefa Biochemia). Seeds were initially sterilized with a solution containing 2% bleach and 0.05% Triton X-100 for 5 min followed by three washes with sterile distilled water and the addition of 0.05% agarose. Plates with seeds were then stratified at 4°C for 2–3 days in the dark. Next, plates were placed in an *in vitro* growth chamber at a temperature of 22°C in a 16-hour light regime. Seedlings were transferred afterwards to soil in a growth chamber with a 16-hour/21°C light and 8-hour/18°C dark regime with 60% humidity. Plants were transformed using the floral dipping method^62^.

### AP-MS on CYCB3;1

Cloning of CYCB3;1 encoding the C-terminal GS^rhino^ tag^63^ fusion under control of the constitutive cauliflower tobacco mosaic virus 35S promoter and transformation of Arabidopsis cell suspension cultures (PSB-D) with direct selection in liquid medium was carried out as previously described^64^.

Pull downs were performed in triplicate, using in-house prepared magnetic IgG beads and 25 mg of total protein extract per pull down as described^64^. On-bead digested samples were analyzed on a Q Exactive (ThermoFisher Scientific) and co-purified proteins were identified with Mascot (Matrix Science) using standard procedures^64^.

After identification, the protein list was filtered versus a large dataset of similar experiments with non-related baits using calculated average Normalized Spectral Abundance Factors (NSAFs)^64^. Proteins identified with at least two matched high confident peptides in at least two experiments, showing high (at least 10-fold) AND significant [-log_10_(*p*-value(T-test)) ≥10] enrichment compared to the large dataset were retained.

### Generation of the CDKB1;1-GFP and GFP-EDE1 reporters

To create the *PRO*_*CDKB1;1*_*:CDKB1;1:EGFP* construct, the genomic fragment of *CDKB1;1* was amplified by PCR and cloned into pDONR221. The SmaI site was inserted in front of the *CDKB1;1* stop codon. *CDKB1;1* constructs were linearized by SmaI digestion and were ligated to the *EGFP* gene, followed by LR recombination reactions with the destination vector *pGWB501*. The same approach was employed to generate the *PRO*_*EDE1*_*:EGFP:EDE1* construct, with the exception that the *EGFP* gene was inserted at the N-terminus of *EDE1* before the first *ATG* codon. Primers used in this study are listed in Table S7.

### Spindle morphogenesis image analysis

First, an ellipse was fitted manually in Fiji to spindles tagged with TagRFP-TUA5 or immunostained against α-tubulin. Next, the major axis, minor axis and spindle area measurements were obtained by going to Analyze > Set measurements and checking the “Area” and “Fit ellipse” boxes. All values are provided in Table S2. To judge the presence of prominent astral microtubules in individual spindle images, spindle files were anonymized in Fiji with the Blind Analysis Tools plugin (https://imagej.net/plugins/blind-analysis-tools). To analyse ψ-tubulin distribution, the images (with a 49 nm pixel size) were first equally treated with the Gaussian Blur filter with a radius of 0.05 scaled units to improve the fluorescence intensity peak definition.

Then, a line was drawn exactly at the middle of the spindle through the pole-to-pole axis in a perpendicular angle in relation to the spindle midzone and the fluorescence intensity profile was plotted in Fiji. The fluorescence intensity values were then normalized by the minimum and maximum values in each cell and combined into a graph containing the mean and SD values of each replicate. The distance between the two highest values of fluorescence was calculated individually in every cell and then corrected by the spindle major axis and plotted as a ratio. In the case of the analysis of GFP-EDE1 distribution, the images (with a 143 nm pixel size) were treated with Gaussian Blur with a radius of 0.1 scaled units.

### Root growth assays and timing of mitotic divisions on oryzalin

For the oryzalin root growth assays, seeds were sown on ½ MS with either 0.05% DMSO as a control or oryzalin. Root growth was recorded daily up until 5 days after germination when plates were scanned and subsequently analyzed with Fiji. To follow mitotic cell divisions on control or oryzalin conditions live, whole five- to seven-day-old seedlings were placed in a glass-bottom dish and covered in solid ½ MS followed by the addition of liquid ½ MS containing 0.05% DMSO as a control or 150 nM oryzalin and incubation for 1 hour. Oryzalin stocks were prepared in DMSO at a concentration of 100 mM and stored at −20°C.

For the root growth assays to assess the functionality of the CDKB1;1-GFP reporter, five-day-old seedlings were transferred onto medium with or without 10 μM Cisplatin for 5 days. At the end of the experiment, plates were photographed and root length was measured using ImageJ software.

### Wholemount immunolocalization of α-tubulin and KNOLLE in roots

Roots of 4-day-old Arabidopsis seedlings were fixed in 4% paraformaldehyde and 0.1% Triton X-100 in MTSB 1/2 buffer (25 mM PIPES, 2.5 mM MgSO4, 2.5 mM EGTA, pH 6.9) for 1 hour under vacuum, then rinsed in PBS 1X for 10 minutes. Samples were then permeabilized in ethanol for 10 minutes and rehydrated in PBS for 10 minutes.

Cell walls were digested using the following buffer for one hour: 2 mM MES pH 5, 0.20% driselase and 0.15% macerozyme. Tissues were hybridized overnight at room temperature with the B-5-1-2 monoclonal anti-α-tubulin (Sigma) and the anti-KNOLLE antibody^65^ (kind gift of G. Jürgens, University of Tubingen, Germany). The next day, tissues were washed for 15 minutes in PBS, 50 mM glycine, incubated with secondary antibodies (Alexa Fluor 555 goat anti-rabbit for KNOLLE antibody and Alexa Fluor 488 goat anti-mouse for the tubulin antibody) overnight and washed again in PBS, 50 mM glycine and DAPI 20 ng/ml. Tissues were mounted in VECTASHIED and DAPI and viewed using an SP8 confocal laser microscope (Leica Microsystems).

Samples were excited sequentially at 405 nm (DAPI), 488 nm (@TUB/Alexa Fluor 488), and 561 nm (@KNOLLE/Alexa Fluor 555), with an emission band of 420-450 nm (DAPI), 495-545 nm (Alexa Fluor 488), and 560-610 nm (Alexa Fluor 555) using a PMT for DAPI imaging, and hybrid detectors for MT and KNOLLE imaging. All stacks were imaged using the same zoom (x 1,60) with a voxel size xyz of 200 nm x 200 nm x 500 nm.

A blind counting was set up to count mitotic MT arrays. Six roots per genotype were analyzed for WT, *cycb3* and *cdkb1*, and seven roots were analyzed for *ede1-1* transformed with GFP-EDE1 WT, GFP-EDE1^8A^ and GFP-EDE1^8D^. All images were first anonymized, and mitotic MT arrays were counted within each root stack using the “Cell counter” ImageJ plugin (https://imagej.nih.gov/ij/plugins/cell-counter.html).

### Immunolocalization of α- and γ-tubulin in root meristematic cells

Root cells were immunostained as described in Liu et al. 1993^66^. α-tubulin was stained using a monoclonal antibody raised in mouse (Sigma, T9026) and γ-tubulin was stained using a monoclonal antibody also raised in mouse (Agrisera, AS20 4482). Since the primary antibodies were raised in the same species, a sequential staining method was employed. First, the slides were incubated with the γ-tubulin antibody overnight at 4°C followed by incubation with the secondary antibody against mouse STAR 635P (abberior) at room temperature for 2 hours. Next, the slides were incubated with the α-tubulin antibody overnight at 4°C followed by incubation with the secondary antibody against mouse STAR 580 (abberior) at room temperature for 2 hours. Samples were then mounted in VECTASHIELD containing DAPI (Vector Laboratories). Slides were imaged in a Zeiss LSM 880 microscope equipped with Airyscan and images were acquired with a voxel size of 49 nm x 49 nm x 160 nm.

### Protein expression and purification and *in vitro* kinase assay

To generate HisGST-EDE1, the *CDS* of EDE1 was initially amplified by PCR with primers containing attB1/attB2 flanking sequences followed by a Gateway BP reaction into pDONR221 and subsequently a Gateway LR reaction into the pHGGWA vector. The destination vector was then transformed in *E. coli* BL21 (DE3) pLysS cells. For expression, *E. coli* cultures were grown until an OD of 0.6 followed by addition of IPTG at a concentration of 0.2 mM and incubation at 16°C overnight. The CDKB1;1-CYCB3;1 complex was expressed and purified as described in Harashima and Schnittger^67^. After purification with Ni-NTA agarose or Strep-Tactin in case of the CDKB1;1 control, all proteins were desalted using PD MiniTrap G-25 columns (GE Healthcare) and protein quality was checked by CBB staining and immunoblotting. Kinase assays were incubated at 30°C for 1 hour in a buffer containing 50 mM Tris-HCl, pH 7.5, 10 mM MgCl_2_, 0.5 mM ATP and 5 mM DTT.

### Sample preparation and LC-MS/MS data acquisition for the identification of EDE1 phosphosites

The protein mixtures were reduced with dithiothreitol, alkylated with chloroacetamide, and digested with trypsin. These digested samples were desalted using StageTips with C18 Empore disk membranes (3 M)^68^, dried in a vacuum evaporator, and dissolved in 2% ACN, 0.1% TFA. Samples were analyzed using an EASY-nLC 1200 (Thermo Fisher) coupled to a Q Exactive Plus mass spectrometer (Thermo Fisher).

For initial assessment of phospho sites, peptides (1:10 dilution) were separated on 16 cm frit-less silica emitters (New Objective, 75 μm inner diameter), packed in-house with reversed-phase ReproSil-Pur C18 AQ 1.9 μm resin (Dr. Maisch). Peptides were loaded on the column and eluted for 50 min using a segmented linear gradient of 5% to 95% solvent B (0 min : 5%B; 0-5 min -> 5%B; 5-25 min -> 20%B; 25-35 min ->35%B; 35-40 min -> 95%B; 40-50 min ->95%B) (solvent A 0% ACN, 0.1% FA; solvent B 80% ACN, 0.1%FA) at a flow rate of 300 nL/min. Mass spectra were acquired in data-dependent acquisition mode with a TOP15 method. MS spectra were acquired in the Orbitrap analyzer with a mass range of 300–1500 m/z at a resolution of 70,000 FWHM and a target value of 3×10^6^ ions. Precursors were selected with an isolation window of 1.3 m/z. HCD fragmentation was performed at a normalized collision energy of 25. MS/MS spectra were acquired with a target value of 5×10^5^ ions at a resolution of 17,500 FWHM, a maximum injection time of 120 ms and a fixed first mass of m/z 100. Peptides with a charge of 1, greater than 6, or with unassigned charge state were excluded from fragmentation for MS^2^; dynamic exclusion for 20s prevented repeated selection of precursors.

For the targeted analysis samples (1:3 dilution) were resolved using the above-mentioned segmented linear gradient. The acquisition method consisted of a full scan method combined with a non-scheduled PRM method. The 17 targeted precursor ions were selected based on the results of DDA peptide search in Skyline. MS spectra were acquired in the Orbitrap analyzer with a mass range of 300–2000 m/z at a resolution of 70,000 FWHM and a target value of 3×10^6^ ions, followed by MS/MS acquisition for the 17 targeted precursors. Precursors were selected with an isolation window of 2.0 m/z. HCD fragmentation was performed at a normalized collision energy of 27. MS/MS spectra were acquired with a target value of 2×10^5^ ions at a resolution of 17,500 FWHM, a maximum injection time of 120 ms and a fixed first mass of m/z 100.

### MS data analysis and PRM method development

Raw data from DDA acquisition were processed using MaxQuant software (version 1.5.7.4, http://www.maxquant.org/)^69^. MS/MS spectra were searched by the Andromeda search engine against a database containing the respective proteins used for the *in vitro* reaction. Trypsin specificity was required and a maximum of two missed cleavages allowed. Minimal peptide length was set to seven amino acids. Carbamidomethylation of cysteine residues was set as fixed, phosphorylation of serine, threonine and tyrosine, oxidation of methionine and protein N-terminal acetylation as variable modifications. The match between runs option was disabled. Peptide-spectrum-matches and proteins were retained if they were below a false discovery rate of 1% in both cases.

Raw data from the DDA acquisition were analyzed on MS1 level using Skyline (https://skyline.ms)^70^ and a database containing the respective proteins used for the *in vitro* reaction. Trypsin specificity was required and a maximum of two missed cleavages allowed. Minimal peptide length was set to seven maximum length to 25 amino acids. Carbamidomethylation of cysteine, phosphorylation of serine, threonine and tyrosine, oxidation of methionine and protein N-terminal acetylation were set as modifications. Results were filtered for precursor charges of 2, 3 and 4. For each phosphorylated precursor ion a respective non-phosphorylated precursor ion was targeted as a control, furthermore several precursor ions from the backbone of EDE1 were chosen as controls between the samples. In total 17 precursors were chosen to be targeted with a PRM approach.

After acquisition of PRM data the raw data were again processed using MaxQuant software, with above-mentioned parameters. Table S6 shows phosphosites and localization probabilities obtained using the MaxQuant search. The mass spectrometry proteomics data have been deposited to the ProteomeXchange Consortium via the PRIDE^71^ partner repository with the dataset identifier PXD046697.

### FRAP assay

For the bleaching of GFP-EDE1, sections of the spindles were bleached with the 405 and 488 lasers both at 100% after 5 frames of imaging and with a scan speed of 7 and 5 iterations. Images were acquired every 0.5 s with a pixel size of 120 nm. For the analysis of the images, the Stowers Plugins Collection was used (https://research.stowers.org/imagejplugins). The data processing and analysis was performed as previously described^72^. For the bleaching of TagRFP-TUA5 in the GFP-EDE1/*ede1-1* and GFP-EDE1^8A^/*ede1-1* backgrounds, only the 405 laser was used at 100% for fluorescence bleaching, but the other parameters were the same as in the bleaching of GFP-EDE1. Outliers in the half maximum values were removed using the ROUT method (Q = 5%).

## Supplemental file legends

**Table S1. Simulation Parameters**

Whenever possible, we used published, experimentally determined values. The configuration file of the simulation is also provided as the definitive source of parameter values.

**Table S2. Quantification and statistical tests of spindle parameters**

**Table S3. Overview of confirmed CYCB3;1 interactors**

A–D: prey annotation. E: number of different bait groups a prey was identified in over the whole AP-MS dataset. Baits were functionally grouped. The lower the more specific. F–G: number of replicates in which a prey was identified with at least two (column F) or with one (column G) unique peptides. H–J: details on the NSAF-based filtering to identify specifically enriched prey proteins.

**Table S4. Protein Identification details obtained with Q Exactive (Thermo Fisher Scientific) and Mascot Distiller software (version 2.5.0, Matrix Science) combined with the Mascot search engine (version 2.6.2, Matrix Science) using the Mascot Daemon interface and database Araport11plus (contaminants filtered out)**

prot_score: protein score; prot_mass: protein mass; prot_matches_sig: number of assigned peptide matches above threshold (high confidence, p < 0.01); prot_sequences_sig: number of significant protein sequences above threshold (high confidence, p < 0.01); prot_cover: percentage of protein sequence covered by assigned peptide matches; prot_len: protein sequence length (AA); prot_pi: pi of identified protein; pep_query: peptide query number; pep_rank: rank of the peptide match, 1 to 10, where 1 is the best match; pep_isbold: peptide is in bold red (Red and bold typefaces are used to highlight the most logical assignment of peptides to proteins. The first time a peptide match to a query appears in the report, it is shown in bold face. Whenever the top-ranking peptide match appears, it is shown in red. Thus, a bold red match is the highest scoring match to a particular query listed under the highest scoring protein containing that match. This means that protein hits with many peptide matches that are both bold and red are the most likely assignments. Conversely, a protein that does not contain any bold red matches is an intersection of proteins listed higher in the report.); pep_isunique: peptide is unique to protein; pep_exp_mz: observed m/z value (precursor); pep_exp_mr: experimental relative molecular mass; pep_exp_z: observed peptide charge state; pep_calc_mr: calculated relative molecular mass; pep_delta: difference (error) between the experimental and calculated masses; pep_start: peptide start position in protein; pep_end: peptide end position in protein; pep_miss: number of missed enzyme cleavage sites; pep_score: peptide ions score; pep_ident: peptide score identity threshold; pep_expect: expectation value for the peptide match (The number of times we would expect to obtain an equal or higher score, purely by chance. The lower this value, the more significant the result); pep_res_before: amino acid before peptide sequence; pep_seq: peptide sequence; pep_res_after: amino acid after peptide sequence; pep_var_mod: any variable modifications found in the peptide; pep_var_mod_pos: position of variable modifications in the peptide.

**Table S5. Mitotic division figures in roots of WT, *cycb3;1 and cdkb1* and GFP-EDE1, GFP-EDE1**^**8A**^ **and GFP-EDE1**^**8D**^ **in the *ede1-1* background**

**Table S6. Phosphorylated sites in EDE1**

**Table S7. Primers used in this study**

